# Disentangling strictly self-serving mutations from win-win mutations in a mutualistic microbial community

**DOI:** 10.1101/530287

**Authors:** Samuel F. M. Hart, Jose Mario Bello Pineda, Chi-Chun Chen, Robin Green, Wenying Shou

**Author notes:** Equal-contribution first author (Samuel F. M. Hart, Jose Mario Bello Pineda).

## Abstract

Mutualisms can be promoted by pleiotropic win-win mutations which directly benefit self (self-serving) and partner (partner-serving). Intuitively, partner-serving phenotype could be quantified as an individual’s benefit supply rate to partners. Here, we demonstrate the inadequacy of this thinking, and propose an alternative. Specifically, we evolved well-mixed mutualistic communities where two engineered yeast strains exchanged essential metabolites lysine and hypoxanthine. Among cells that consumed lysine and released hypoxanthine, a chromosome duplication mutation seemed win-win: it improved cell’s affinity for lysine, and increased hypoxanthine release rate per cell. However, increased release rate was due to increased cell size accompanied by increased lysine utilization per birth. Consequently, total hypoxanthine release rate per lysine utilization (defined as “exchange ratio”) remained unchanged. Indeed, this mutation did not increase partner or community growth rate, and is thus solely self-serving. By extension, reduced benefit production rate by an individual may not imply cheating.

## Introduction

Mutualisms, mutually beneficial interactions between species, are widely observed between microbes (Goldford et al., 2017; Morris et al., 2013, 2012; Seth and Taga, 2014) and between microbes and their hosts (Seth and Taga, 2014). Often, mutualisms involve the release and consumption of essential metabolites such as vitamins and amino acids (Beliaev et al., 2014; Carini et al., 2014; Helliwell et al., 2011; Jiang et al., 2018; Rodionova et al., 2015; Zengler and Zaramela, 2018). Extensive metabolic interactions between microbes have been thought to contribute to the difficulty of culturing microbes in isolation (Kaeberlein et al., 2002). Under certain conditions, microbial metabolic exchanges may even promote community growth (Tasoff et al., 2015).

In mutualistic communities, a mutation can exert instantaneous effects on the individual itself as well as on the mutualistic partner. For example, a mutation could increase self’s growth rate by reducing benefit supply to a mutualistic partner. Although the mutant will harm itself in the long term as its partner suffers, we would classify this mutation as “self-serving, partner-harming” (“win-lose”) instead of “self-harming, partner-harming” (“lose-lose”). Qualitatively, the instantaneous fitness effect of a mutation on self and on partner can be positive, neutral, or negative, giving rise to 3×3=9 types.

Distinguishing mutation types is important for predicting their evolutionary successes. Consider the general case of microbial mutualisms without any partner choice mechanisms. That is, an individual is not capable of discriminating or “choosing” among spatially-equivalent partners (Sachs et al., 2004; Shou, 2015). Then, a well-mixed environment will favor mutations with a positive instantaneous fitness effect on self (i.e. selfish, strictly self-serving, and win-win; Fig 1A, “mixed”). This is because in a well-mixed environment, benefits from mutualistic partners are uniformly distributed, and thus how much an individual contributes to mutualistic partners is irrelevant. In contrast, in a spatially-structured environment, mutations exerting a positive instantaneous effect on the mutualistic partner (i.e. win-win, strictly partner-serving, and altruistic) can be favored, while selfish mutations can be disfavored (Chao and Levin, 1981; Doebeli and Knowlton, 1998; W. D. Hamilton, 1964; Harcombe, 2010; Momeni et al., 2013b; Nowak, 2006; Sachs et al., 2004; Shou, 2015) (Fig 1A, “spatial”). This is because in a spatially-structured environment, interactions are localized and repeated between neighbors. If an individual does not aid its mutualistic neighbor, the individual will eventually suffer as its mutualistic neighbor perishes. Win-win mutations are particularly intriguing because they directly promote both sides of a mutualism.

**Fig 1.**
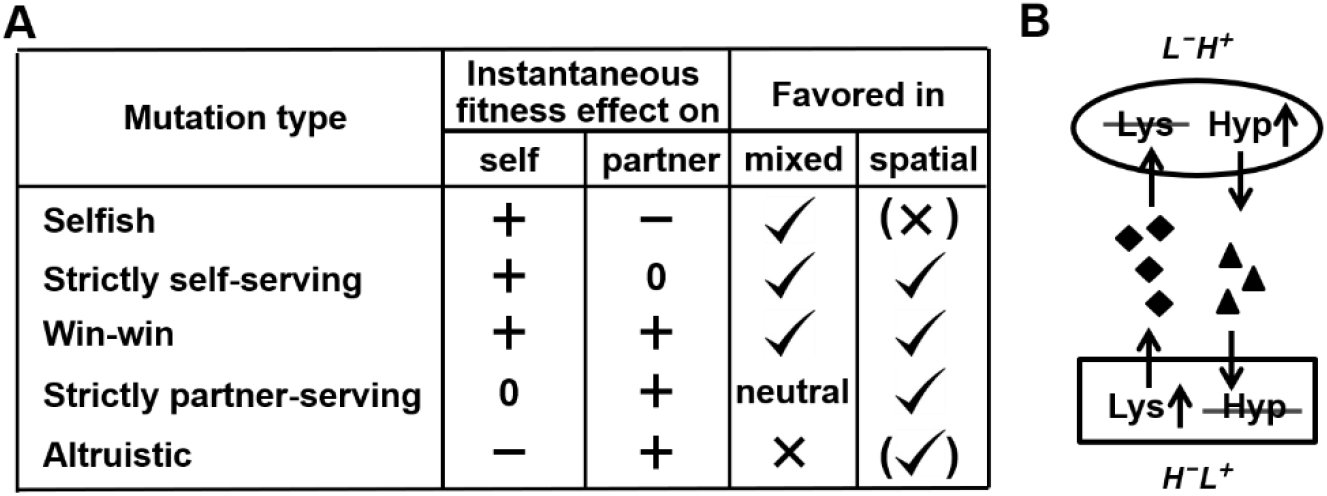
Mutation types predicted to be favored during the evolution of a mutualistic community. **(A) Mutation types in a mutualistic community and their evolutionary fates.** Mutations that exert a positive instantaneous effect on self (selfish, strictly self-serving, and win-win) are favored in a well-mixed environment. In a spatially-structured environment, effects on self and on partner are both important. For example, a spatially-structured environment may favor an altruistic mutation that confers a large benefit on partner at a small cost to self. Parentheses indicate that selection outcome (favored or disfavored) depends on quantitative details of the fitness effects on self and partner (see (Momeni et al., 2013b) for an example). **(B) CoSMO.** CoSMO is an engineered mutualistic community consisting of two non-mating *S. cerevisiae* strains (Hart et al., 2019a; Shou et al., 2007). Thus, the two strains may be regarded as two species. The mCherry-expressing *L*^−^*H*^+^ strain is unable to synthesize lysine (*L*) and overproduces the adenine precursor hypoxanthine (*H*). The complementary GFP-expressing *H*^−^*L*^+^ strain requires hypoxanthine and overproduces lysine. Both overproduction mutations render the first enzyme of the corresponding biosynthesis pathway insensitive to end-product feedback inhibition control (Armitt and Woods, 1970; Feller et al., 1999). In minimal medium lacking exogenously supplied *L* and *H*, the two strains form a mutualistic community where live cells from both strains release overproduced metabolites (Hart et al., 2019a) and support each other’s growth.

Here, we analyze a mutation arising during the evolution of an engineered yeast mutualistic community in a well-mixed environment. This community CoSMO (<u>C</u>ooperation that is <u>S</u>ynthetic and <u>M</u>utually <u>O</u>bligatory) (Shou et al., 2007) consists of two *S. cerevisiae* strains that interact via metabolite cross-feeding (Figure 1B). *L*^−^ *H*^+^ requires lysine (*L*) and overproduces and releases the adenine precursor hypoxanthine (*H*) (Hart et al., 2019a). The complementary *H*^−^*L*^+^ requires *H*, and overproduces and releases *L*. The two yeast strains are reproductively isolated, and thus can be regarded as two species. This mutualism is “cooperative” in the sense that metabolite over-production is costly to both strains (Figure 1-Figure Supplement 1; (Waite and Shou, 2012)). Mutualisms modeled by CoSMO are widely observed in natural communities (Beliaev et al., 2014; Carini et al., 2014; Helliwell et al., 2011; Jiang et al., 2018; Rodionova et al., 2015; Zengler and Zaramela, 2018), including those in the gut and oral microbiota (Palmer et al., 2001; Rakoff-Nahoum et al., 2014). Indeed, principles learned from CoSMO, including how fitness effects of species interactions affect the composition and spatial patterning of member species, and mechanisms that protect mutualisms from exploiters, have been found to operate in communities of non-engineered microbes (Momeni et al., 2013a, 2013b; Waite and Shou, 2012). CoSMO offers an ideal system for examining the evolution of mutualism, given the genetic tractability of the budding yeast and the validated phenotype quantification methods (Hart et al., 2019a).

In this study, we demonstrate that a seemingly intuitive definition of partner-serving phenotype can lead to erroneous conclusions. We will conclude by discussing how to quantify important theoretical concepts such as “benefit”, “cost”, and “partner-serving phenotype”, especially for microbial mutualisms where interactions span multiple generations.

## Results

We propagated nine independent CoSMO communities in well-mixed minimal medium without lysine or hypoxanthine supplements for ~100 generations by performing periodic dilutions to keep culture turbidity below saturation (Methods, “CoSMO evolution”). We froze samples at various time points for later revival. Since all communities were well-mixed, we predicted that selfish, strictly self-serving, and win-win mutations should arise (Fig 1A). In this study, we focused on *L*^−^*H*^+^.

### Evolved L^−^H^+^ clones harboring Chromosome 14 duplication appeared to display a win-win phenotype

We randomly isolated evolved *L*^−^*H*^+^ clones from independent communities. Since the community environment was lysine-limited (Hart et al., 2019a; Waite and Shou, 2012), improved growth under lysine limitation would be self-serving. Indeed, while the ancestral strain failed to grow into micro-colonies on agar with low lysine (1.5 μM), all tested (>20) evolved clones could (Fig 2-Figure Supplement 1; Methods “Microcolony assay”), consistent with our previous findings (Hart et al., 2019a; Waite and Shou, 2012). Thus, evolved *L*^−^*H*^+^ clones displayed self-serving phenotypes.

Since hypoxanthine is also scarce in the community (Hart et al., 2019a), a mutant *L*^−^*H*^+^ cell with increased hypoxanthine release rate should allow the partner to grow faster and is thus partner-serving. We randomly chose several evolved *L*^−^*H*^+^ clones, and measured their hypoxanthine release rate (Hart et al., 2019a) (Methods, “Release assay”; Fig 3-Figure Supplement 1). Whereas two of the evolved clones released hypoxanthine at a similar rate as the ancestor, three increased release rates in the sense that each cell released more hypoxanthine per hour than the ancestor (Fig 3-Figure Supplement 2). We then sequenced genomes of both types of clones (Methods, “Whole-genome sequencing”; Fig 2-Figure Supplement 2). Chromosome 14 duplication (*DISOMY14*) occurred in all three clones that exhibited faster-than-ancestor hypoxanthine release rate (Fig 3 - Figure Supplement 2, blue), and not in the other two clones that exhibited ancestral release rate (Fig 3-Figure Supplement 2, orange).

When we back-crossed evolved clones harboring *DISOMY14* to the ancestral background (Methods), only meiotic segregants containing *DISOMY14* showed increased hypoxanthine release rate compared to the ancestor (Fig 3 – Figure Supplement 3). Thus, *DISOMY14* genetically co-segregated with increased release rate (partner-serving). *DISOMY14* repeatedly rose to a detectable frequency when *L*^−^*H*^+^ evolved with *H*^−^*L*^+^ in CoSMO (3 out of 3 lines), or when *L*^−^*H*^+^ evolved alone in lysine-limited chemostats (5 out of 5 lines). Thus, *DISOMY14* is likely adaptive in lysine limitation. Indeed, unlike ancestors, meiotic segregants containing *DISOMY14* could form micro-colonies on low-lysine plate (self-serving). Taken together, we hypothesized *DISOMY14* to be both self-serving and partner-serving, i.e. “win-win”.

### The self-serving phenotype of DISOMY14 requires duplication of the lysine permease LYP1

Chromosome 14 harbors the high-affinity lysine permease *LYP1*. To test whether *LYP1* duplication might improve the growth rate of *L*^−^*H*^+^ in limited lysine, we inserted an extra copy of *LYP1* into the ancestral *L*^−^*H*^+^ strain (Methods, “Gene knock-in and knock-out”), and quantified cell growth rate under various concentrations of lysine using a microscopy assay (Methods, “Microscopy growth assay”) (Hart et al., 2019b). *LYP1* duplication indeed significantly increased the growth rate of *L*^−^*H*^+^ in low lysine (Fig 2, compare green with magenta). Similarly, deleting the duplicated copy of *LYP1* from *DISOMY14* cells abolished the self-serving phenotype of *DISOMY14* (Fig 2, compare blue and cyan with magenta). Taken together, duplication of *LYP1* is responsible for the self-serving phenotype of *DISOMY14*.

**Fig 2.**
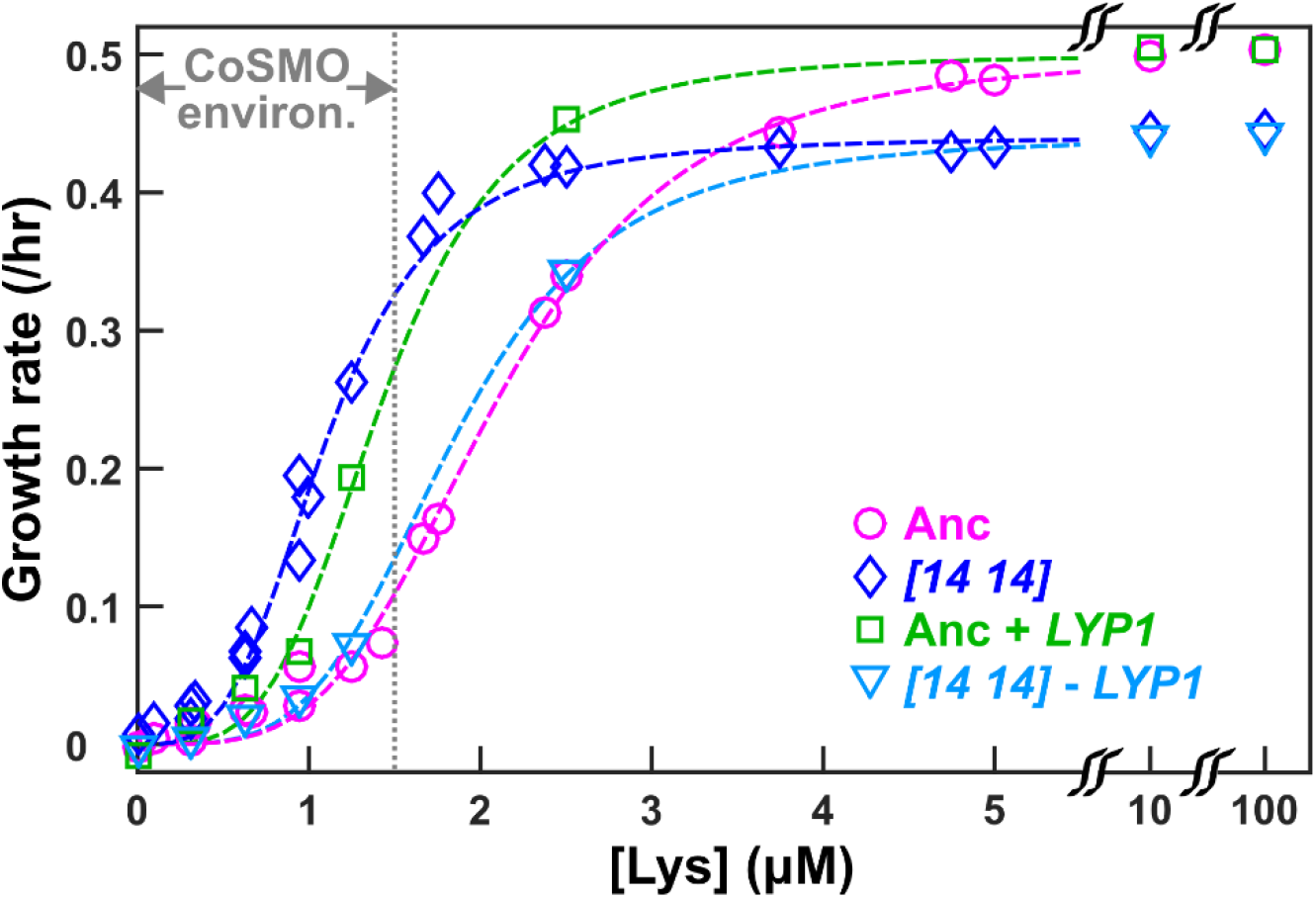
DISOMY14 improves L^−^H^+^ growth rate under lysine limitation via duplication of the high-affinity lysine permease LYP1. Exponentially-growing cells were washed free of lysine, starved in SD for 3~5 hrs to deplete vacuolar lysine storage, and incubated in microtiter wells containing SD supplemented with various concentrations of lysine. Cells were imaged using a fluorescence microscope, and total fluorescence was tracked over time (Methods; “Microscopy growth assay”) (Hart et al., 2019b). The maximal growth rate (the steepest positive slope of ln[fluorescence] against time) was quantified and plotted against lysine concentration. The grey dotted line demarcating “CoSMO environ.” corresponds to the lysine level supporting a growth rate of ≤ 0.1/hr as observed in ancestral CoSMO. *DISOMY14* (“*[14 14]*”, blue diamond; WY2261) grew faster than the ancestral *L*^−^*H*^+^ (magenta circle; WY1335) in low lysine. This self-serving phenotype of *DISOMY14* was abolished when the duplicated *LYP1* gene was deleted form *DISOMY14* (cyan triangle; WY2262, WY2263). Conversely, introducing an extra copy of *LYP1* into the ancestor improved grow rate in limited lysine (green square; WY2254, WY2255). Each data point is the average of multiple (~4) experiments. Dashed fitting curves: Moser growth equation *g* = *g_max_L^n^*/(*K^n^* + *L^n^*) where *g* is growth rate of *L*^−^*H*^+^, *L* is lysine concentration, *g_max_* is the maximal growth rate in excess lysine, *K* is the lysine concentration supporting *g_max_*/2, and *n* is the “growth cooperativity” constant describing the sigmoidal shape of the curve. The maximal growth rate of *DISOMY14* is lower than that of the ancestor, which could be due to the fitness cost associated with aneuploidy (Oromendia et al., 2012). Data can be found in Fig 2 Source Data.

### WHI3 duplication is responsible for the increased hypoxanthine release rate of DISOMY14

To identify which duplicated gene(s) might be responsible for the increased hypoxanthine release rate of *DISOMY14*, we systematically deleted various sections of the duplicated Chromosome 14 (Fig 3A; Methods, “Chromosome truncation”), and quantified hypoxanthine release rate. Duplication of the region between *YNL193W* and *GCR2* was necessary for the increased release rate (Fig 3B, orange). This region contains six genes, including *WHI3*. Integrating an extra copy of *WHI3* into the ancestor increased release rate to near that of *DISOMY14*, while deleting one copy of *WHI3* from *DISOMY14* restored the ancestral release rate (Fig 4A; Methods “Gene knock-in and knock-out”).

**Fig 3.**
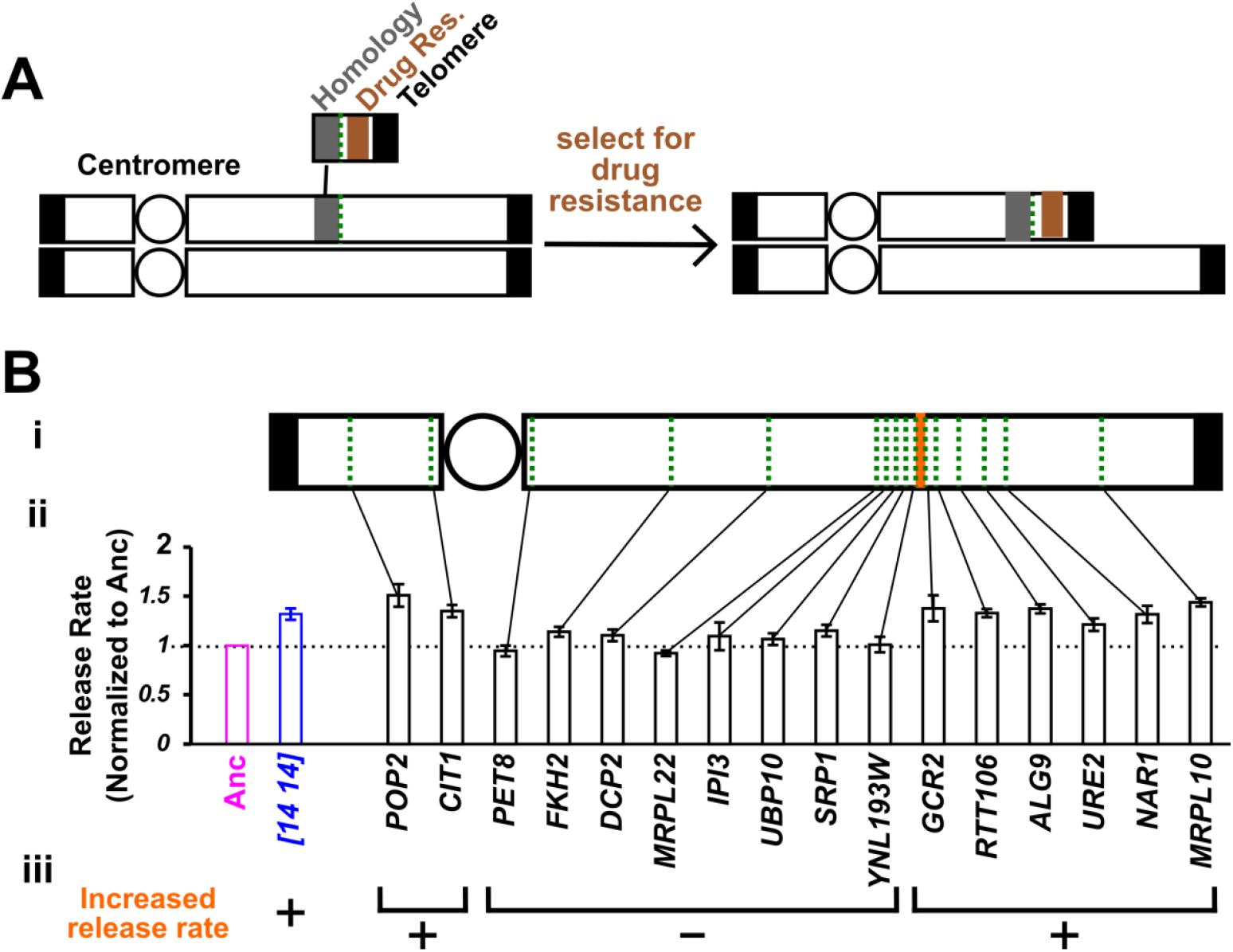
Chromosome truncation reveals the regions responsible for increased lysine affinity and increased hypoxanthine release rate. **(A) Chromosome truncation scheme.** Recombination occurs between a chromosome and a truncation cassette containing a homology region, a drug resistance marker, and a telomere (Methods, “Chromosome truncation”). Single integration leads to the truncation of one of the duplicated chromosomes, with the centromere (circle)-containing region being retained by the cell. Integration into both chromosomes would lead to an inviable cell due to loss of the segment immediately distal to the insertion site from both chromosomes (not drawn). The copy number of chromosomal regions in transformants was verified by RADseq (Methods “RADseq”; Fig 3-Figure Supplement 4). **(B) The genomic region between *YNL193W* and *GCR2* is responsible for increased release rate.** We systematically truncated chromosome 14 in *DISOMY14* cells (WY2261). We chose truncation sites (green dotted lines) that were spread across chromosome 14, and in later rounds of truncation, spread across the region of interest (i). We quantified hypoxanthine release rates of transformants in starvation batch cultures (Methods “Release assay”; Fig 3-Figure Supplement 1), normalized them against the release rate of the ancestral strain measured in the same experiment, and plotted the mean value and two standard errors of mean (ii). Duplication of the region between *YNL193W* and *GCR2* (orange; containing six genes including *YNL193W* and excluding *GCR2*) is responsible for the increased hypoxanthine release rate. Data can be found in Fig 3 Source Data.

**Fig 4.**
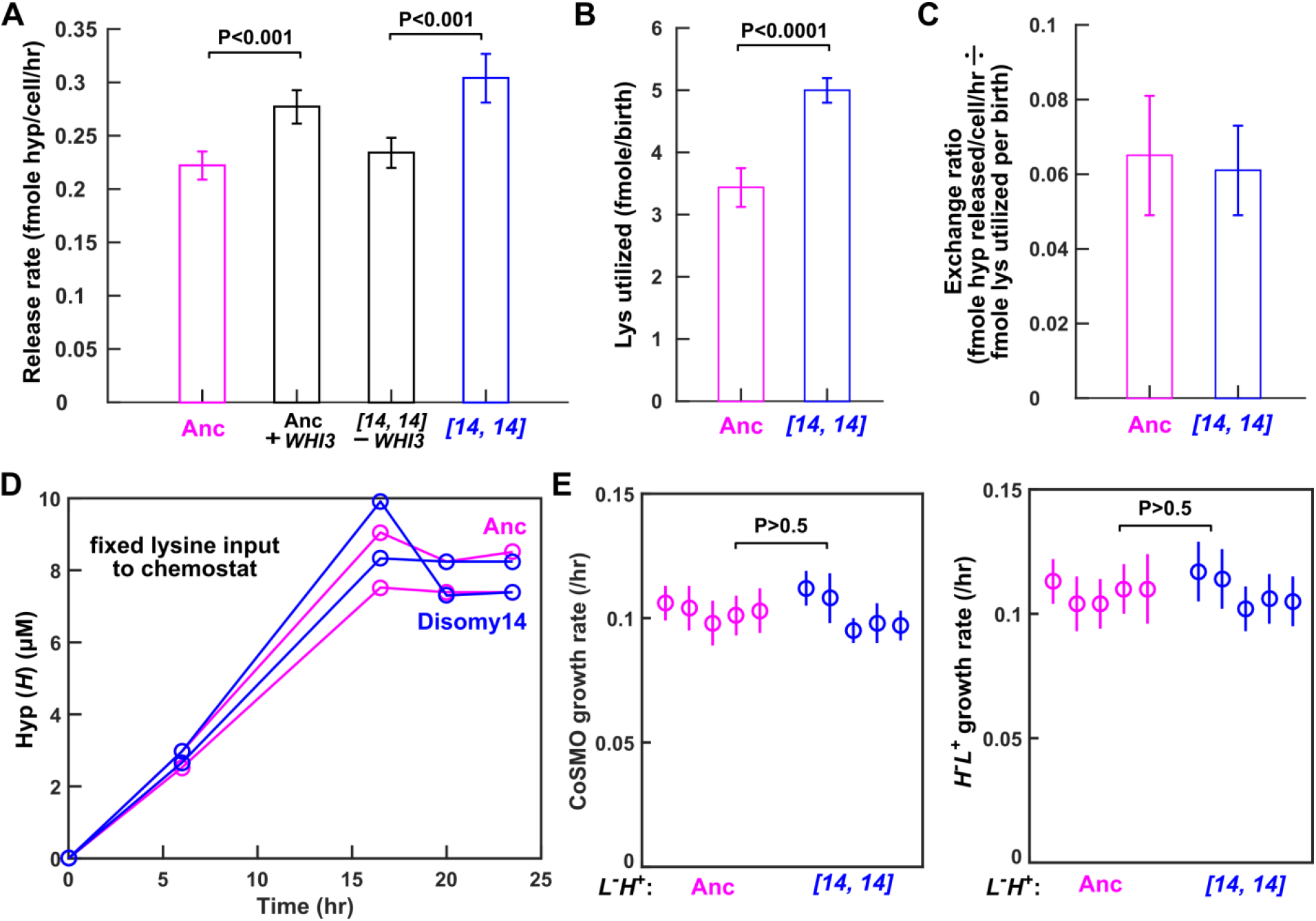
Despite increased release rate per cell, DISOMY14 is not partner-serving. **(A) *WHI3* duplication is responsible for increased hypoxanthine release rate per cell**. Introducing an extra copy of *WHI3* into the ancestral background (WY2357~2359) increased hypoxanthine release rate per cell. Deleting the duplicated *WHI3* from *DISOMY14* (WY2350~2352) decreased release rate. **(B) *DISOMY14* utilizes more lysine per birth than the ancestor.** For A and B, mean and two SEM are plotted. **(C) Indistinguishable exchange ratio between ancestor and *DISOMY14*.** Hypoxanthine release rate per *L*^−^*H*^+^ cell (A) is divided by lysine utilization per *L*^−^*H*^+^ birth (B). Error bars are calculated via error propagation (Methods). A-C were measured in starvation batch cultures. **(D) In chemostats fed with a fixed lysine supply rate, *DISOMY14* and ancestor release the same total hypoxanthine.** Ancestor (WY1335, magenta) or *DISOMY14* (WY2349, blue) *L*^−^ *H*^+^ cells were grown in lysine-limited chemostats with a doubling time of 6 hrs (Methods “chemostats”). Supernatant hypoxanthine concentrations were quantified using a yield-based bioassay (Methods, “Bioassay”; Fig 3-Figure Supplement 4). For complete data and statistical comparison, see Fig 4 Figure Supplement 2. **(E) *DISOMY14* and ancestral *L*^−^ *H*^+^ led to identical growth rate of community and of partner.** To prevent rapid evolution in *L*^−^*H*^+^ (Hart et al., 2019a), we grew CoSMO in a spatially-structured environment on agarose pads, and periodically measured the absolute abundance of the two strains (differentiable by their fluorescence; Methods, ‘flow cytometry’). We then quantified the steady-state growth rate of community and of partner *H*^−^*L*^+^ by regressing ln(cell density) against time after the initial lag phase up to <10^8^ cells ((Hart et al., 2019a); see Fig 4 - Figure Supplement 3 for examples of detailed dynamics). We plotted the slope (i.e. growth rate), with error bars indicating 2x standard error of estimating the slope. In A, B, and E, we performed statistical comparisons first using the F-test to test for equal variance, and then using unpaired two-tailed *t*-test with equal variance. We plotted the corresponding P-values of the *t*-test (the probability of observing a test statistic as extreme as, or more extreme than, the observed value under the null hypothesis of two groups belonging to the same distribution). Comparisons in A and B show significant difference, while those in E are not significantly different. All data can be found in Fig 4 Source Data.

### Despite increased release rate per cell, DISOMY14 is not partner-serving

*WHI3* encodes a cell cycle inhibitor that binds the mRNA of *CLN3* (an activator of cell cycle) (Garí et al., 2001; Nash et al., 2001). Deletion of *WHI3* is known to result in smaller cell size, whereas extra copies or overexpression of *WHI3* is known to increase cell size (Garí et al., 2001; Nash et al., 2001). Indeed, *DISOMY14* cells are bigger than ancestral cells, as quantified by the Coulter counter (Fig 4-Figure Supplement 1A; Methods “Cell size measurements”). Integrating an extra copy of *WHI3* into the ancestor increased mean cell size, while deleting the extra copy of *WHI3* from *DISOMY14* restored ancestral cell size (Fig 4-Figure Supplement 1A).

Consistent with *DISOMY14* cells being bigger than ancestral cells, *DISOMY14* cells utilized more lysine per birth than the ancestor (Fig 4B; Methods “Metabolite utilization in batch culture”). Integrating an extra copy of *WHI3* into the ancestor increased lysine utilization per newborn cell, while deleting the extra copy of *WHI3* from *DISOMY14* reduced lysine utilization per newborn cell (Fig 4-Figure Supplement 1B). Thus, although each *DISOMY14* cell released more hypoxanthine, each also utilized more lysine.

When we normalized hypoxanthine release rate per *L*^−^*H*^+^ cell (*r_H_*) by lysine utilization per *L*^−^*H*^+^ birth (*u_L_*), which we define as the “*H-L* exchange ratio” (*r_H_/u_L_*), *DISOMY14* is indistinguishable from the ancestor (Fig 4C). In other words, a fixed amount of lysine can be converted to ancestral *L*^−^*H*^+^ cells which are more numerous but lower-releasing (Fig 5B, i), or *DISOMY14* cells which are fewer but higher-releasing (Fig 5B, ii). As long as the ancestor and *DISOMY14* displayed a similar *total* hypoxanthine release rate per lysine utilized, the partner *H*^−^*L*^+^ would not benefit more from one than the other. In Discussions, we will describe how exchange ratio links to “inclusive fitness” or “direct fitness”.

**Fig 5.**
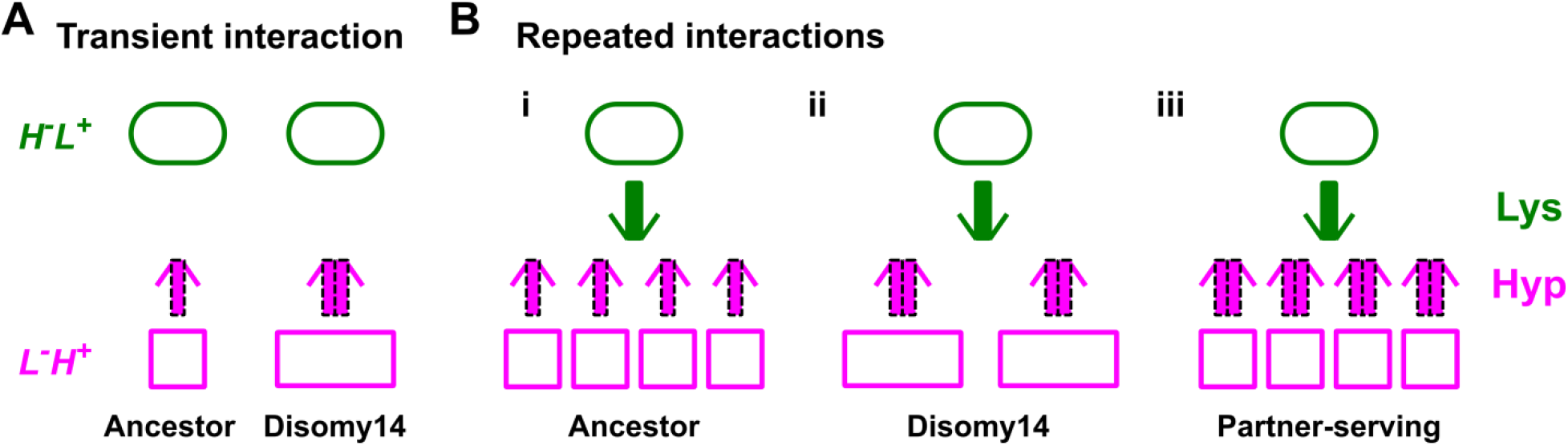
Schematic illustration of partner-serving mutations. (**A**) Transient interaction. At the initial stage of an interaction, *DISOMY14* with an increased release rate per cell (2x thick purple arrow) is more partner-serving than the ancestor (1x thick purple arrow), consistent with the biological market theory. (**B**) Long-term repeated interactions. Over multiple generations, *DISOMY14* is not partner-serving. Even though each *DISOMY14* cell releases more hypoxanthine than each ancestral cell, a fixed amount of lysine (green arrow) is turned into more ancestral cells than *DISOMY14* cells (compare i and ii). A true partner-serving mutation should increase total benefit supply rate per intake benefit (iii), which is equivalent to (benefit supply rate per cell)/(benefit utilized to make the cell).

If *DISOMY14* is not partner serving, then fixed lysine supply would lead *DISOMY14* and ancestor to release identical total hypoxanthine. To test this hypothesis, we cultured ancestral and *DISOMY14 L*^−^*H*^+^ cells in lysine-limited chemostats with a fixed rate of lysine supply (Methods, “Chemostat”). Indeed, the steady state hypoxanthine concentrations in chemostats were indistinguishable between ancestor and *DISOMY14* (Fig 4D). Similar to batch culture results, in chemostats *DISOMY14* released hypoxanthine at a higher rate than the ancestor on a per cell basis, but utilized more lysine per birth, leading to an identical *H-L* exchange ratio as the ancestor (Fig 4 – Figure Supplement 2B). Consistent with *DISOMY14* not being partner serving, partner *H*^−^*L*^+^ grew at the same rate whether co-cultured with ancestral or *DISOMY14 L*^−^*H*^+^ (Fig 4E, right panel).

Taken together, despite having a higher hypoxanthine release rate per cell, *DISOMY14* is not partner-serving or win-win. Rather, *DISOMY14* is strictly self-serving.

## Discussions

Interpreting social phenotypes can be surprisingly nuanced (Dubravcic et al., 2014; Gore et al., 2009; Greig and Travisano, 2004; Kümmerli and Ross-Gillespie, 2014; Rainey et al., 2014; Smith et al., 2013; Zhang and Rainey, 2013). Here, by dissecting the molecular bases of how *DISOMY14* affects self (*L*^−^*H*^+^) and partner (*H*^−^*L*^+^), we illustrate how microbial win-win or partner-serving phenotypes should be interpreted. The self-serving aspect of *DISOMY14* is straightforward: due to duplication of the lysine permease gene *LYP1, DISOMY14* grows faster than the ancestor in lysine limitation typically found in the CoSMO environment (Fig 2). However, the partner-serving aspect of *DISOMY14* is far less straightforward: we had mis-interpreted *DISOMY14’*s increased hypoxanthine release rate per cell as partner-serving until we realized that duplication of *WHI3*, a cell cycle inhibitor which made cells larger, was responsible. We advocate that when considering microbial mutualisms that span multiple generations, a focal individual’s partner-serving phenotype may be quantified as its exchange ratio (total benefit release rate per metabolite utilized, or equivalently, benefit release rate per cell divided by metabolite utilization per birth). Below, we discuss how exchange ratio links to current frameworks of mutualisms, inclusive fitness theory, and biological market theory.

### Theories of mutualisms

General theories have been developed for mutualisms (Archetti et al., 2011; Doebeli and Knowlton, 1998; Foster and Wenseleers, 2006; Frank, 1994; Jones et al., 2015; Sachs et al., 2004; Trivers, 1971; West et al., 2002; Yamamura et al., 2004). In mathematical models of mutualisms (e.g. (Doebeli and Knowlton, 1998; Foster and Wenseleers, 2006; Frank, 1994; Yamamura et al., 2004)), exchanged goods (investments) were linked into fitness effects on the focal individual. For example, the net fitness gain of the focal individual = fitness gain per investment made * average investment made within the group – fitness loss per investment made* investment by the focal individual (Frank, 1994). For a focal *H*^−^*L*^+^, the fitness loss term (fitness loss per lysine released*total lysine released) is fixed whether the interaction partner is ancestral or *DISOMY14 L*^−^ *H*^+^. The term of fitness gain per investment made (how much faster *H*^−^ *L*^+^ would grow per unit of lysine supplied to *L*^−^*H*^+^) embodies the spirit of exchange ratio (total hypoxanthine release rate per unit of lysine), although measuring fitness gain per investment can be difficult since it is not a constant.

In exchange ratio, “benefit” and “investment” are defined in physical terms of the goods exchanged, and fitness “cost” for making the investment is separately considered (see Eq. 1 below). Specifically, when interacting with a focal *L*^−^*H*^+^ cell, the benefit and investment by *H*^−^*L*^+^ can be defined from two equivalent perspectives. In the “individualistic” perspective, we can define the benefit gained by *H*^−^*L*^+^ as the release rate of hypoxanthine by the focal *L*^−^*H*^+^ cell (*r_H_*), while investment made by *H*^−^*L*^+^ as the amount of lysine required to make that *L*^−^*H*^+^ cell (*u_L_*).

Then this ratio of benefit-to-investment for *H*^−^*L*^+^ is identical to the exchange ratio for *L*^−^ *H*^+^. On the other hand, in the “population” perspective, we can define the investment made by *H*^−^*L*^+^ as a unit of lysine released for *L*^−^ *H*^+^, and the benefit received by *H*^−^*L*^+^ as the resultant *total* rate of hypoxanthine reciprocation from not only the focal *L*^−^*H*^+^ cell but also its offspring (Fig 5B). This benefit-to-investment ratio is also identical to the exchange ratio, since total hypoxanthine release rate/fmole lysine = (hypoxanthine release rate per cell * total number of *L*^−^*H*^+^ cells)/fmole lysine = *r_H_*/(fmole lysine/total number of *L*^−^*H*^+^ cells)= *r_H_/u_L_*.

### Inclusive fitness

Why would an individual cooperate - paying a fitness cost to provide a benefit that can aid the reproduction of other individuals (e.g. sterile ant workers aiding the reproduction of the queen)? Social evolution theories offer explanations for the evolution of cooperative traits (Frank, 1998; W. D. Hamilton, 1964; Kerr, 2009; Lehmann and Keller, 2006; Maynard Smith, 1964; Price, 1972, 1970; Queller, 1985; Sachs et al., 2004; Traulsen and Nowak, 2006; West et al., 2007).

A central concept in social evolution theories is “inclusive fitness” where the fitness impact of social interactions is considered and where natural selection leads organisms to become adapted as if to maximize their inclusive fitness (Grafen, 2006). Hamilton’s rule states that cooperation can evolve as long as *rb-c*>0, where *c* is the fitness cost to the focal cooperator, *b* is the benefit to its partner, and *r* is their “relatedness” – the similarity of an actor to its recipient relative to the population (Damore and Gore, 2012; Fletcher and Doebeli, 2009, 2009; W.D. Hamilton, 1964; Queller, 1992; van Veelen, 2009). Note that “similarity” can broadly refer to action type (e.g. cooperation versus no cooperation), even if the actor and the recipient are genetically unrelated as in mutualisms. A mathematically-equivalent, individual-centric version of inclusive fitness (also known as “direct fitness”) of a focal individual is the sum of its basal fitness *w_0_* in an asocial environment plus *rb* benefit received from other cooperators in the social environment minus the cost of cooperation *c* (Damore and Gore, 2012; Foster and Wenseleers, 2006; Frank, 1998; West et al., 2007). If a focal individual’s inclusive fitness is greater than its basal fitness (*w_0_*+*rb-c*> *w_0_* or *rb-c*>0), then the individual would grow faster by cooperating than by not cooperating. Inclusive fitness relies on additivity of fitness effects and other assumptions. However in microbial communities, fitness often increases in a nonlinear (e.g. sigmoidal or saturating) fashion as the benefit increases, and therefore inclusive fitness described above is thought to be over-simplified and sometimes even misleading (Damore and Gore, 2012; Grafen, 2006; Nowak et al., 2010; van Veelen, 2009). However, despite these criticisms, inclusive fitness has served as a useful conceptual framework (Abbot et al., 2011; Birch Jonathan, 2017).

In obligatory mutualisms between clonal populations (i.e. relatedness =1), the two partners’ exchange ratios and costs of making investments together generate a physical definition of inclusive fitness or direct fitness. This fitness term turns out to be identical between the two partners, which equals the entire community’s growth rate. Specifically, the basal fitness of *H*^−^*L*^+^ and *L*^−^ *H*^+^ in monoculture is negative due to death rate and the cost of making investments. When growing in communities, divergent strain ratios rapidly converge to a fixed value (Momeni et al., 2013a; Shou et al., 2007). With their ratio fixed, the two strains must grow at an identical rate equal to the community growth rate *g_comm_*. In other words, the growth rate of the two strains are identical to each other and to *g_comm_*:

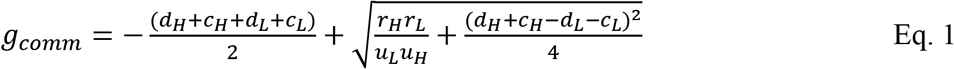

((Hart et al., 2019a), Methods “Community growth rate”), where *d_H_* and *d_L_* are respectively *H*^−^*L*^+^ and *L*^−^*H*^+^’s death rates, *c_H_* and *c_L_* are respectively *H*^−^*L*^+^ and *L*^−^*H*^+^’s fitness costs of overproducing metabolites, *r_H_* and *r_L_* are respectively hypoxanthine and lysine release rates, and *u_H_* and *u_L_* are respectively hypoxanthine and lysine utilization amount per birth. Note that 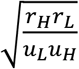 is the geometric mean of the two exchange ratios *r_H_/u_L_* and *r_L_/u_H_*, and *c_H_* and *c_L_* contain the fitness costs of making investments for the partner.

A corollary to Eq. 1 is that despite *DISOMY14*’s better affinity for metabolite (Fig 2), *DISOMY14* does not enjoy improved inclusive fitness *g_comm_* compared to the ancestor when cultured with partner *H*^−^*L*^+^ (Fig 4E). This is because in CoSMO, the growth of an individual is limited by partner-supplied metabolites. In other words, there is no point of eating faster if meals arrive slowly. Obviously, if we were to culture *DISOMY14 L*^−^*H*^+^, ancestral *L*^−^*H*^+^, and *H*^−^*L*^+^ in a well-mixed environment, then *DISOMY14* should outcompete ancestor due to its better affinity for lysine (Fig 2). This also brings up an important point: the exchange ratio-based fitness metric works for cases where each mutualistic partner is of one genotype. It does not capture changes in genotype frequencies within a species.

We might apply this exchange ratio-based fitness metric to sub-communities in a spatially-structured environment. In a spatially-structured environment, if the initial population densities are low, then interactions will mainly take place between one genotype from each species, and individuals in the local sub-community with the highest inclusive or direct fitness are predicted to grow the fastest. In the case of spatial CoSMO, would an *H*^−^*L*^+^ cell grow faster if it had landed next to an ancestral or a *DISOMY14 L*^−^*H*^+^ cell? During the initial encounter, *H*^−^*L*^+^ next to *DISOMY14* will benefit more from *DISOMY14*’s faster release rate (Fig 5A). However, after this initial stage, *DISOMY14* and ancestor are identical, since similar total hypoxanthine release rate is generated per fmole of utilized lysine (Fig 5B, i and ii). Indeed, *H*^−^*L*^+^ grew equally fast when co-cultured with ancestral or *DISOMY14 L*^−^*H*^+^ (Fig 4E).

### Biological market theory

Biological market theory posits that the exchange of goods among organisms can be analyzed in market terms, where individuals attempt to maximize their gains (Noe and Hammerstein, 1994; Noë and Hammerstein, 1995; Werner et al., 2014). For example, a male insect that offers more nuptial gifts to a female is regarded by the female as being more partner (female)-serving and is thus chosen by the female (Noe and Hammerstein, 1994; Noë and Hammerstein, 1995; Werner et al., 2014). Although biological market theory does not apply here since our yeast strains lack partner choice capability, our work can prove useful for other systems. For example, consider the legume-rhizobia mutualism where a legume host provides photosynthates to rhizobia while rhizobia reciprocate with fixed nitrogen. Application of biological market theory leads to statements such as “…whether plant hosts can detect variation in resources or services provided (by rhizobia) and respond accordingly. Such discrimination mechanisms have been found in legumes, with some species preferentially supporting rhizobial symbionts that provide more fixed N_2_ for hosts” (Werner et al., 2014). Since legume-rhizobia mutualisms last over multiple generations of rhizobia, we suggest that “fixed N_2_” should be quantified in terms of the exchange ratio: the total release rate of fixed N_2_ per photosynthate utilized, or a focal rhizobium’s nitrogen release rate normalized by photosynthate utilized to make the rhizobium. Obviously, when comparing non-nitrogen fixers with nitrogen fixers, fixers are more mutualistic than non-fixers and thus making the fine distinction of exchange ratio is unnecessary. But when comparing quantitative variants of nitrogen fixers, this distinction could become important. By explicitly quantifying the exchanged goods, exchange ratio captures the spirit of market economy.

In summary, when considering microbial mutualisms that span multiple generations, a focal individual’s partner-serving phenotype can be quantified as the exchange ratio of benefit production rate for partner by focal cell divided by benefit utilized to make the focal cell. This is equivalent to total benefit production rate for partner per intake benefit. By the same token, an individual with reduced benefit production rate may not be a cheater if the individual also utilizes less benefit from the partner.

## Methods

### Strains and medium

Genetic manipulations and growth medium for the yeast *S. cerevisiae* are explained in (Guthrie and Fink, 1991). Protocols and technical details that we have used can be found in (Waite and Shou, 2014). Briefly, we used rich medium YPD (10 g/L yeast extract, 20 g/L peptone, 20 g/L glucose) in 2% agar plates for isolating single colonies. Saturated YPD overnight liquid cultures from these colonies were then used as inocula to grow exponential cultures. We sterilized YPD media by autoclaving. YPD overnights were stored at room temperature for no more than 4~5 days prior to experiments.

We used defined minimal medium SD (6.7 g/L Difco™ yeast nitrogen base w/o amino acids, 20 g/L glucose) for all experiments (Guthrie and Fink, 1991), with supplemental metabolites as noted (Waite and Shou, 2014). To achieve higher reproducibility, we sterilized SD media by filtering through 0.22 μm filters. To make SD plates, we autoclaved 20 g/L Bacto™ agar or agarose in H_2_O, and after autoclaving, supplemented equal volume of sterile-filtered 2XSD.

The ancestral strain was WY1335, described in detail in (Hart et al., 2019a). All strains are in Table S1.

The *DISOMY14* strain we used for analysis WY2261 (refrozen as WY2348 and WY2349) was obtained in the following manner. The evolved strain WY1584 was back-crossed twice into the ancestral background to get rid of mutations in *ECM21* and *YPL247C*. The first cross with WY1521 resulted in “38-1D”, which was then crossed with WY1335 to result in WY2261 (“E2”). To genotype spores, we PCR amplified the mutated regions in *ECM21* and *YPL247C*, and subjected the purified PCR product to Sanger sequencing. For those spores that contained no mutations in *ECM21* and *YPL247C*, we subjected them to restriction-site associated DNA sequencing (RADseq; Methods) to determine ploidy. When we modified our sequence analysis pipeline, we realized that WY2261 contained other mutations (Table S2). However, the presence of other mutations does not affect our conclusions, since integrating an extra copy of *LYP1* or *WHI3* into the ancestral background respectively increased growth rate under lysine limitation (Figure 2) and per cell hypoxanthine release rate (Figure 4A).

### CoSMO evolution

*L*^−^*H*^+^ (WY1335) and *H*^−^*L*^+^ (WY1340) were grown separately to exponential phase in minimal SD medium supplemented with lysine (164.3 μM) or adenine sulfate (108.6 μM), respectively (Guthrie and Fink, 1991). Cells were washed free of supplements, counted using a Coulter counter, and mixed at 1000:1 (Line A), 1:1 (Line B), or 1:1000 (Line C) at a total density of 5×10^5^/ml. The different initio ratios did not noticeably affect evolutionary outcomes. Three 3ml community replicates (replicates 1, 2, and 3) per initial ratio were initiated. Communities were grown at 30°C in glass tubes on a rotator to ensure well-mixing. Community turbidity was tracked by measuring the optical density (OD_600_) in a spectrophotometer once to twice every day. In this study, 1 OD was found to be 2~4×10^7^cells/ml. We diluted communities periodically to maintain OD at below 0.5 to avoid additional selections due to limitations of nutrients other than adenine or lysine. The fold-dilution was controlled to within 10~20 folds to minimize introducing severe population bottlenecks. Note that no mutagens were used during evolution.

Coculture generation was calculated from accumulative population density by multiplying OD with total fold-dilutions. For each coculture at every 10~20 generations, the cell pellet of ~1ml coculture was resuspended in 1ml rich medium YPD (Guthrie and Fink, 1991) + 10% trehalose, cooled at 4°C for several hours, and frozen at −80 °C. Cells frozen this way revived much better than if frozen in SD medium supplemented with a final of 15% glycerol.

CoSMO could engage in self-sustained growth only if its initial total cell density was sufficiently high (Shou et al., 2007). Thus, to revive a coculture, ~20 μl was scooped from the frozen stock using a sterile metal spatula, diluted ~10-fold into SD, and allowed to grow to moderate turbidity. The coculture was further expanded by adding 3ml of SD. To isolate clones, cocultures were plated on rich medium YPD and clones from the two strains were distinguished by their fluorescence colors or drug resistance markers.

### Gibson Assembly

The detailed protocol of Gibson assembly (Gibson et al., 2009) for assembling DNA fragments with end homology was obtained from Eric Klavins lab (UW). 1ml 5xISO buffer included: 1M Tris-HCl (pH 7.5) 500 μl; 2M MgCl2 25μl; 100 mM dGTP, dATP, dCTP, and dTTP 10μl each (total 40 μl); 1M DTT 50μl; 100mM NAD 50μl; PEG-8000 0.25g; H_2_O: 145 μl. 5xISO was frozen in 100 μl aliquots at −20°C.

The assembly master mix (375 μl total) included: H_2_O 216.75μl; 5XISO Buffer 100μl; 1 U/μl T5 Exonuclease 2μl; 2U/μl Phusion Polymerase 6.25μl; 40U/μl Taq DNA Ligase 50μl. 15 μl aliquots were stored at −20°C. This master mix is ideal for DNA molecules with 20~150 bp overlapping homology. To carry out the assembly reaction, 15 μl assembly master mix is mixed with a total of 5μl DNA (e.g. 125 nM), and incubated at 50°C for 1hr.

### Chromosome truncation

We constructed plasmid WSB175 to contain G418 resistance (*KanMX*) and a telomeric sequence. Briefly, WSB174 (from Dan Gottschling lab) containing a *URA3* marker was digested with HindIII and BamHI to remove the *URA3* marker and yield the vector backbone (4.5kb). The *KanMX* gene was amplified from WSB26 using primers WSO433 and WSO434, each containing a 25-bp overhang homologous to the vector backbone. The vector backbone and the *KanMX* PCR product were circularized via Gibson assembly to yield WSB175. To perform Gibson assembly, we used 5 μl DNA (including 55ng vector and 0.5 μl of 125 nM insert). Gibson mixture was transformed into *E. coli*, and DNA was extracted from several colonies. DNA was checked via restriction digestion with HindIII and BamHI.

To carry out chromosome truncation, we PCR amplified ~600 bp fragments from various genomic locations on Chromosome 14 (Fig 3B). The PCR reaction consisted of: genomic DNA (0.5 μl out of 25 μl where 0.3 ml overnight yeast culture was harvested and DNA extracted), 10xPCR buffer (5 μl), 25 mM MgCl_2_ (3 μl), 10mM dNTP mix (1 μl), 50 μM forward primer (0.5 μl), 50 μM reverse primer (0.5 μl), Taq polymerase (0.5 μl), H_2_O (39 μl). Cycling conditions are as following: 94°C 3 min; [94°C 30 sec + 56.9°C 60 sec + 72°C 60 sec] x30 cycles; 72°C 10 min. We used Qubit (Thermofisher) to quantify DNA concentration. We then assembled PCR fragment with vector backbone containing *KanMX* and telomere (2.5 μl of each at 125 nM, corresponding to ~100ng insert and 350ng vector) to yield the assembled DNA. The assembled DNA (100ng) was PCR amplified again. PCR reaction contained: Gibson product (4.1 μl), 5xPhusion HF buffer (10 μl), 10mM dNTP mix (1 μl), 50 μM forward primer (1 μl), 50 μM reverse primer (WSO157, 1 μl), Phusion polymerase (0.5 μl), H_2_O (32.4 μl). Cycling conditions are as following: 98°C 30 sec; [98°C 10 sec + 59.9°C 30 sec + 72°C 90 sec] x30 cycles; 72°C 10 min. Expected length was ~2.4kb.

The PCR product was used to transform the *DISOMY14* yeast strain (WY2261) using lithium acetate yeast transformation. Transformants were selected on rich medium supplemented with G418 (YPD+G418) plate. Transformants were screened for correct integration using PCR amplification across the chromosome integration site (one primer homologous to the genome, and the other primer homologous to *KanMX*).

### Gene knock-in and knock-out

To introduce an extra copy of *LYP1* in the *ste3::HygMX* locus of the ancestor, we assembled and transformed the following. We amplified a 527 bp homology region upstream of *STE3*, the *LYP1* gene (including 333 bp upstream and 466 bp downstream of ORF), and the *KanMX* resistance cassette (loxP-TEFp-KanMX-TEFt-loxP) from WSB118, all using PCR. Primers used contained 20-bp homology when appropriate to allow us to compile these sequences in the order listed using Gibson assembly. We then amplified this assembly further using PCR and transformed this 4.8 kB product into the ancestor (WY1335), screening for successful integration by G418 resistance, loss of hygromycin resistance, and checking PCR. We obtained WY2254~2255.

To knockout duplicated *LYP1* from *DISOMY14*, we amplified a 532bp region upstream of *LYP1*, and a 739bp region downstream of *LYP1*. We also amplified KanMX resistance. We assembled the three pieces via Gibson assembly, PCR amplified the assembled molecule, and transformed *DISOMY14* cells with the PCR product. Transformants were plated on YPD+G418 plate, and colonies were screened for correct integration via PCR. We obtained WY2262~2263.

We used a similar methodology to introduce an extra copy of *WHI3* in the *ste3::HygMX* locus of the ancestor (WY1335). We first digested WSB185 (pBSKII) with XmaI and HindIII (HF), yielding 2.9 kb backbone. We amplified a 428bp homology region upstream of *STE3*, the *WHI3* gene (including 570bp upstream and 700bp downstream of ORF; ~3.3kb), and *KanMX* resistance (1.6kb), and assembled all these with the vector backbone. The Gibson product was used to transform *E. coli*, and colonies were mini-preped and screened via restriction digest with Kpn1 and BamH1. The correct liberated fragment (5.4kb) was transformed into ancestral cells (WY1335) and plated on YPD+G418 plate. Transformants were checked for loss of hygromycin resistance and via PCR. We obtained WY2357~2359.

To delete *WHI3* from *DISOMY14*, we amplified a 528bp region upstream of *WHI3*, and a 489bp region downstream of *WHI3*. We assembled the two pieces with KanMX cassette via Gibson assembly, PCR amplified the assembled molecule, and transformed *DISOMY14* cells with the PCR product. Transformants were plated on YPD+G418 plate, and colonies were screened for correct integration via PCR. We obtained WY2350~2352.

### Genomic DNA extraction for sequencing

For whole-genome sequencing using tagmentation, we have extracted yeast genomic DNA used QIAGEN Genomic-tip 20G (Cat. No. 10223), YeaStar Genomic DNA kit (Zymo Research), and a protocol modified from Sergey Kryazhimskiy and Andrew Murray lab. DNA from the last protocol is suitable for tagmentation but not for RADseq.

To extract DNA, we used the following procedure: Spin down cells (0.5ml saturated culture; microfuge at highest speed in 2ml v-bottom tubes for 2’). Thoroughly discard supernatant. To the pellet add 252 μl Digestion mix (50μl of 0.5M EDTA, pH = 7.5 or 8.0; 200μl of ddH20; 2.5μl of Zymolyase (stock: 5U/ul)). Mix by inversion and incubate at 37°C for 24h on rotator. Add 50μl of miniprep mix (0.2M EDTA, 0.4M Tris, 2% SDS, pH = 8.0). Mix by inversion and incubate at 65°C for 30min. Add 63μl (about 1/5 of volume) of 5M KAc. Mix by inversion and incubate on ice for 30 min. Add 250μl of chloroform, vortex vigorously for 1min. This helps to precipitate proteins and lipids. Spin down sample for 10min on max speed (tabletop centrifuge). The DNA is in the supernatant. Transfer supernatant to a new tube (max 300μl). Add 720μl (> 2x volume) of 100% EtOH. Mix by inversion. At this point you should see the DNA clots in your tube. Spin down on max speed for 20 minutes. The DNA is now in the pellet. Thoroughly discard supernatant. Add 50μl H_2_0 + 1μl of RNAase A (10mg/ml) to undried pellet. Allow DNA to resuspend by incubating at 37°C for 1hr. Add 2μl of Proteinase K (20mg/ml). Incubate for 2hrs at 37°C. Add 130μl of 100% isopropanol. Mix by inversion and spin down at max speed for 10min. The DNA is in the pellet. Thoroughly discard supernatant. Add 500μl of 70% EtOH. Mix by inversion and spin down at max speed for 10min. The DNA is in the pellet. Discard supernatant. Allow pellet to air dry overnight. Resuspend pellet in 100μl of 10mM Tris, pH = 8.0.

For RADseq, yeast genomic DNA was extracted from 2×10^8^~10^9^ cells using, for example, the DNeasy Blood & Tissue Kit (Qiagen). High-quality DNA is required for optimal restriction endonuclease digestion and is of utmost importance for the overall success of the protocol. The samples were treated with RNase A following manufacturer’s instructions to remove residual RNA then quantified using Qubit. The optimal concentration after elution is 25 ng/μl or greater.

### Whole-genome sequencing

The whole genome sequencing protocol was slightly modified from that of Sergey Kryazhimskiy v. 2.1 (2013-06-06) (Kryazhimskiy et al., 2014). Indexing primer design followed (Adey et al., 2010) (Table S3). For an illustration of Nextera V2 Illumina sequencing molecular biology, see Fig 2-Figure Supplement 2.

To tagment genomic DNA, we prepared gDNA at concentration at or below 2.5ng/μl. For *n* samples = *r* rows and *c* columns, make the Tagmentation Master Mix (TMM) by mixing *n* x 1.06 x 1.25μl of TD Buffer (Tagment DNA Buffer) and *n* x 1.06 x 0.25μl of TDE1 (Tagment DNA Enzyme) in a PCR tube. Mix thoroughly by gently pipetting the mixture up and down 20 times. Distribute TMM into *r* tubes (or a PCR strip), *c* x 1.03 x 1.5μl into each tube. With a multichannel pipette, distribute TMM into all wells of a fresh plate (“tagmentation plate”), 1.5μl per well. With a multichannel pipette, transfer 1μl of gDNA into the tagmentation plate (total volume = 2.5μl per well). Mix by gently pipetting up and down 10 times. Cover plate with Microseal ‘B’(Biorad, MSB-1001). Give the plate a quick spin to collect all liquid at the bottom (Sorvall or Allegra centrifuges, 1000 rpm for 1 min). Place the plate in the thermocycler and run the following program: 55°C for 5 min; hold at 10°C.

Next, PCR amplification is performed to add the index adaptors to tagmented DNA. We will make the adaptor PCR reaction final volume to be 7.5 μl (2.5 μl of tagmented DNA from above, 3.75 μl of 2x KAPA master mix (KAPA amplification kit KK2611/KK2612), 0.625 μl of Index Adapter 1, 0.625 μl of Index Adapter 2). For convenience, we have pre-mixed index primers where index adapter 1 is always the same (NexV2ad1noBC; Table S3) and index adapter 2 is one of the 96 Index adaptors (NexV2ad2**; Table S3), and each is at 5 μM in H_2_O. Mix the entire mix by gently pipetting up and down 10 times. Cover plate with Microseal ‘A’ (Biorad, MSB-5001). Make sure to press well on each well, especially edge wells. Give the plate a quick spin to collect all liquid at the bottom at 1000 rpm for 1 min. Place the tubes in the thermocycler and run the following program: 72°C for 3 min; 98°C for 2:45 min; [98°C for 15 sec; 62°C for 30 sec; 72°C for 1:30 min]x8; Hold at 4°C. Ensure that the lid is tight and that it is heated during incubation.

Make Reconditioning PCR Master Mix (RMM) by mixing *n*\*1.06*8.5 μl of 2xKAPA polymerase mix, *n*\*1.06*0.5 μl of primer P1 (10μM; WSO380 AATGATACGGCGACCACCGA), and *n*\*1.06*0.5 μl of primer P2 (10μM; WSO381 CAAGCAGAAGACGGCATACGA). Mix thoroughly by gently pipetting the mixture up and down 20 times. With a multichannel pipette, transfer 9.5 μl of RMM into each well of the plate (final PCR volume 17μl). Mix by gently pipetting up and down 10 times. Cover plate with Microseal ‘A’. Give the plate a quick spin to collect all liquid at the bottom at 1000 rpm for 1 min. Place the tubes in the thermocycler and run the following program: 95°C for 5 min; [98°C for 20 sec; 62°C for 20 sec; 72°C for 30 sec]x4; 72°C for 2 min; Hold at 4°C.

PCR clean-up used magnetic beads. Centrifuge the plate to collect all liquid (1000 rpm for 1 min). Vortex AMPure XP beads (Beckman Coulter A63880) for 30 sec to ensure that they are evenly dispersed. Transfer *c* x 1.05 x 1 x *V* μl (*V*=PCR volume=17μl) of beads into *r* PCR tubes or a PCR strip. Using a multichannel pipette, transfer *V* μl of beads into each well containing the PCR product. Mix well by gently pipetting up and down 20 times. The color of the mixture should appear homogeneous after mixing. Incubate at room temperature for 5 min so that DNA is captured by beads. Place the plate on the magnetic stand (Life Technologies, Cat. #123-31D) and incubate for about 1 min to separate beads from solution. Wait for the solution to become clear. While the plate is on the magnetic stand, aspirate clear solution from the plate and discard. Do not disturb the beads. If beads are accidentally pipetted, resuspend them back, wait for the solution to clear up, and repeat. While the plate is on the magnetic stand, dispense 200μl of 70% ethanol into each well and incubate for 30 seconds at room temperature. Aspirate out ethanol without disturbing the beads and discard. Repeat for a total of 2 washes. Remove the remaining ethanol with P10 pipette. Let the plate air dry for approximately 5 min. Do not overdry the beads. Take the plate off the magnetic stand. Add 33μl of 10mM Tris-HCl (pH 8) to each well of the plate. Carefully resuspend the beads by mixing 10-15 times. Incubate for 2 min at room temperature. DNA is now in the solution. Place the plate back onto the magnetic stand and incubate for about 1 min to separate beads from solution. Wait for the solution to become clear. While the plate is on the magnetic stand, aspirate clear solution from the plate and transfer to a fresh plate. Do not disturb the beads. If beads are accidentally pipetted, resuspend them back, wait for the solution to clear up, and repeat. Qubit quantify samples using 1 μl of eluate. We get about 6ng/μl. Send 3 μl for High Sensivitity tapestation (75-1000 pg/ul) to get average length. If Qubit reading was >0.38ng/μl, then sequencing worked.

Pool samples at equal molarity, with the final pool should ideally be at least 2 nM (although 1 nM seemed fine as well). If we have 100 indexes, then each sample needs to be diluted to 0.02 nM. Qubit the pooled sample and submit 30 μl at 2nM to the sequencing facility, and sequenced on Illumina HiSeq 2000 (paired-end; 50~150 cycles; Nextera sequencing primers).

### RADseq

RADseq (restriction site-associated DNA sequencing) protocol was obtained from Aimee Dudley lab based on (Etter et al., 2012). The design scheme is in Fig 3-Figure Supplement 5, and primer sequences are in Table S4. Briefly, genomic DNA was digested with the six-cutter Mfe1 and the four-cutter Mbo1. Our desired DNA fragment would be flanked by Mfe1 and Mbo1 sites. To these, we ligate annealed primers (P1 top annealed with P1 bottom containing a 4bp barcode and Mfe1 overhang, and P2 top annealed with P2 bottom containing a 6-bp barcode and Mbo1 overhang). The dual barcode system allows many samples (e.g. 900 samples) to be sequenced simultaneously. P1 also contains Illumina Read 1 sequencing primer which will read 4bp barcode and genomic DNA adjacent to the Mfe1 site, and P2 also contains Illumina Read 2 sequencing primer and Index sequencing primer which will read 6bp barcode and genomic DNA adjacent to the Mbo1 site.

The ligation product can be PCR amplified using WSO381 and WSO398 (Fig 2-Figure Supplement 2). P2 top primer has a stretch of sequences (lower case) that does not anneal with P2 bottom. Thus, in PCR round 1, only WSO398 is effective. In PCR round 2 or later, both WSO398 and WSO381 are effective. Note that the non-annealing sequence is designed for the following reason: Because most genomic DNA fragments are flanked by the Mbo1 site and these fragments would ligate with P2 on both sides, the non-annealing DNA segment (lower case) ensures that these fragments lacking P1 cannot be amplified with WSO381.

To anneal P1 and P2, 100 μM stock plates of P1 (top and bottom) and P2 (top and bottom) primers were obtained from the Dudley Lab at the Pacific Northwest Research Institute. The P1 bottom primer and the P2 top primer include a 5’-phosphate modification required by ligation. For the P1 annealing reaction, the following was mixed: 10 μl 5M NaCl, 100 μl 1M Tris (pH 8.0), 888 μl H_2_O, and 1 μl each 100 μM P1 top and P1 bottom primers (final concentration 100 nM). For the P2 annealing reaction, the following was mixed: 7 μl 5M NaCl, 70 μl 1M Tris (pH 8.0), 483 μl H_2_O, and 70 μl each 100 μM P1 top and P1 bottom primers (final concentration 10 μM). The P1 and P2 top/bottom primer mixtures were aliquoted into a PCR tubes at 100 μl per tube. The samples were heated at 95°C for 1 min, then allowed to cool to 4°C (at a rate of 0.1°C/sec) to allow annealing of the top and bottom primers. After cooling, the tubes were spun down and immediately put on ice. The P1 aliquots were consolidated into a single tube, and P2 aliquots were consolidated into a single tube.

The P1 and P2 annealed primers were combined as follows. 700 μl of P2 was diluted into 4.55 ml of H_2_O (diluted concentration at 1.33 μM). To 48 wells of a 96 well plate, 83 μl of this diluted annealed P2 to each well. 27.5 μl of the annealed P1 primers was added to each well. The final P1+P2 volume is 110.5 μl (P2 final concentration 1μM; P1 final concentration of 25 nM). It is very important to keep adapters cold after annealing. Specifically, keep at 4°C, mix on ice, thaw on ice after retrieving from −20°C storage.

RNase-treated yeast genomic DNA was subjected to restriction digestion by Mfe1 and Mbo1 (NEB). Per reaction, 5.25 μl of the purified genomic DNA (~125 ng, Qubit quantified) was mixed with 0.625 μl 10X NEB Buffer 4 (or CutSmart buffer), 0.25 μl MboI (2.5 units), and 0.125 μl MfeI-HF (2.5 units) in PCR tubes (final volume of 6.25 ul). The mixture was set up on ice and incubated at 37°C for 1 hour, and was heat inactivated by incubating at 65°C for 20 minutes.

Custom adapters were annealed to the digested DNA using NEB T4 DNA ligase. Specifically, the following components were added to a tube in this order: 2.5 μl of the combined annealed P1 (25 nM)+P2 (1 μM) adaptor mix, the entire 6.25 μl digestion reaction, and 3.75 μl of a T4 ligase master mix (0.1 μl T4 ligase at 2000 units/μl, 1.25 μl T4 ligation buffer, and 2.4 μl H_2_O). The reaction was combined on ice and mixed by pipetting up and down. Ligation was carried out at room temperature for 20 minutes and heat inactivated in thermocycler at 65°C for 20 minutes. The samples were allowed to cool slowly to room temperature (30 min).

Every 24 samples were pooled together and concentrated using the QIAGEN MinElute PCR purification kit, and the MinElute column was eluted in 10 μl EB. The concentrated samples were subjected to gel electrophoresis using 2% low range ultra agarose gel (48 samples/lane). Sufficient band separation is achieved when the loading dye is approximately halfway down the gel, which ensures that the 150 band is separated from the 100 bp primer dimer. The size range 150 to 500 bp was excised out of the gel under a long-wavelength UV lamp and purified using the QIAGEN MinElute Gel Extraction Kit, eluting with 20 μl EB.

The gel extracted DNA was amplified using the NEBNext PCR Master Mix (NEB#M0541S) with custom primers (WSO398 and WSO381; Table S4). The mixture contained: 25 μl NEBNext PCR Master Mix; 1 μl WSO398 (10μM); 1 μl WSO381 (10μM); 1 μl DNA (5~10ng), and 22 μl H_2_O (total 50 μl). PCR cycling was: 98 °C (1 min); [98 °C (10 sec) + 60 °C (30 sec) + 72 °C (30 sec)]x14 cycles + 72 °C (4 min) + 4 °C hold. The PCR reaction was cleaned and concentrated using the QIAquick PCR Purification Kit (QIAGEN), eluting in 30 μl H_2_O. The expected concentration is ~30-40 ng/μl. The library quality (fragment size distribution) was ascertained using Tapestation. The resultant DNA was subjected to paired-end 25 cycles on Illumina HiSeq 2000 using TruSeq Dual Index Sequencing Primers).

### Sequence analysis

To analyze whole genome sequencing, a custom Perl script incorporating bwa (Li and Durbin, 2009) and SAMtools (Li et al., 2009) written by Robin Green was used to align paired-end reads to the *S. cerevisiae* RM-11 reference genome. Mutations were identified via GATK for single-nucleotide variants and indels, and cn.MOPs for local copy-number variant calling. A custom Perl script incorporating vcftools was used to automate evolved versus ancestral strain comparison. All genetic changes were visually inspected using the Integrated Genome Viewer (IGV) environment for quality inspection and validation. Ploidy was calculated using custom python and R scripts wherein read depth was counted for each base. These read depths were averaged within successive 1000-bp windows; each window average is normalized by the median of all window averages across the genome. The normalized values for each window are log2 transformed and plotted versus the respective genomic position (chromosome/supercontig) for ease in graphical inspection of ploidy changes. Sequence analysis code can be publicly accessed at https://github.com/robingreen525/ShouLab_NGS_CloneSeq.

RADseq analysis was performed as described in (Etter et al., 2012) using custom python and R scripts. Samples were split by their respective barcode and aligned to the RM11 reference genome using bwa. Up to 6 mismatches were allowed per read/marker. Next, reads with Phred quality scores below 20 or with a median coverage of less than 2 per sample were discarded. To ensure that each respective marker was representative of a properly digested MfeI-MboI, the expected length of each fragment based on a theoretical digestion of the RM11 genome was compared to the length of the actual marker as determined by read alignment to that marker (i.e finding reads that fell within the expected coordinates of a MfeI-MboI digest product). Next, for each marker, the proportion of reads aligning to that marker was normalized against total read alignment to the genome.

To ensure that only high quality markers were used, the CV of each marker across all tested strains were analyzed and markers with a CV of >=0.6 were discarded. Additionally, only markers that were within the expected gel cut size of >125 bp and <400 bp were used. This still allowed >2000 markers to be used for downstream analysis.

To assess ploidy for Radseq, the same analysis was performed on a panel of 10 euploid strains. For each strain and for each marker, the relative proportion of that maker of the total reads for the strain of interest was compared against the median proportion of the total reads for the euploid panel. A supercontig (the RM11 assembly does not have full chromosomes but supercontigs) was called as duplicated if the average proportion of all makers on that supercontig in the backcrossed strain was 2-fold greater than the euploid panel. All disomy 14 calls for a tetrad segregated 2:2 as expected.

### Microcolony assay

This method has been described in (Hart et al., 2019b). Briefly, to assay for self-serving phenotype of an *L*^−^*H*^+^ mutant, we diluted a saturated overnight 1:6000 into SD+164 μM lysine, and allowed cultures to grow overnight at 30°C to exponential phase. We washed cells 3x with SD, starved them for 4-6 hours to deplete vacuolar lysine stores, and diluted each culture so that a 50 μl spot had several hundred cells. We spotted 50 μl on SD plate supplemented with 1.5 μM lysine (10 spots/plate), and allowed these plates to grow overnight. When observed under a 10x objective microscope, evolved cells with increased lysine affinity would grow into “microcolonies” of ~20~100 cells, while the ancestral genotype failed to grow (Fig 2-Figure Supplement 1). *DISOMY14* exhibited an intermediate phenotype where smaller microcolonies with variable sizes formed.

### Flow cytometry

Beads (ThermoFisher Cat R0300, 3 μm red fluorescent beads) were autoclaved in a factory-clean glass tube, diluted into sterile 0.9% NaCl, and supplemented with sterile-filtered Triton X-100 to a final 0.05% (to prevent bead clumping). The mixture was sonicated to eliminate bead clusters and was kept at 4°C in constant rotation to prevent settling and re-clumping. Bead density was quantified via hemacytometer and Coulter counting (4-8×10^6^ beads/ml final). The prepared bead mixture served as a density standard. Culture samples of interest were diluted to OD 0.01~0.1 (7×10^5^ - 7×10^6^ cells/ml) in filtered water. Bead-cell mixtures were prepared by mixing 90 ul of the diluted culture sample, 10 μl of the bead stock, and 2 μl of 1 μM ToPro 3 (Molecular Probes T-3605), a nucleic acid dye that only permeates cell membranes of dead cells. Triplicate cell-bead mixtures were prepared for each culture in a 96-well format for high-throughput processing. Flow cytometry of the samples was performed on Cytek DxP Cytometer equipped with four lasers, ten detectors, and an autosampler. mCherry and ToPro are respectively detected by 75 mW 561nm Laser with 615/25 detector and 25mW 637nm laser with 660/20 detector. Each sample was individually analyzed using FlowJo^®^ software to identify the number of beads, dead cells, and live fluorescent cells. Live and dead cell densities were calculated from the respective cell:bead ratios, corrected for the initial culture dilution factor. The mean cell density from triplicate measurements was used (coefficient of variation within 5% to 10%).

### Microscopy growth assay

See (Hart et al., 2019b) for details on microscopy and experimental setup, method validation, and data analysis. Briefly, cells were diluted to low densities to minimize metabolite depletion during measurements. Dilutions were estimated from culture OD measurement to result in 1000~5000 cells inoculated in 300 μl SD medium supplemented with different metabolite concentrations in wells of a transparent flat-bottom microtiter plate (e.g. Costar 3370). We filled the outermost wells with water to reduce evaporation.

Microtiter plates were imaged periodically (every 0.5~2 hrs) under a 10x objective in a Nikon Eclipse TE-2000U inverted fluorescence microscope. The microscope was connected to a cooled CCD camera for fluorescence and transmitted light imaging. The microscope was enclosed in a temperature-controlled chamber set to 30°C. The microscope was equipped with motorized stages to allow z-autofocusing and systematic xy-scanning of locations in microplate wells, as well as motorized switchable filter cubes capable of detecting a variety of fluorophores. Image acquisition was done with an in-house LabVIEW program, incorporating bright-field autofocusing (Hart et al., 2019b) and automatic exposure adjustment during fluorescence imaging to avoid saturation. Condensation on the plate lid sometimes interfered with autofocusing. Thus, we added a transparent “lid warmer” on top of our plate lid (Hart et al., 2019b), and set it to be 0.5°C warmer than the plate bottom, which eliminated condensation. We used an ET DsRed filter cube (Exciter: ET545/30x, Emitter: ET620/60m, Dichroic: T570LP) for mCherry-expressing strains.

Time-lapse images were analyzed using an ImageJ plugin Bioact (Hart et al., 2019b). Bioact measured the total fluorescence intensity of all cells in an image frame after subtracting the background fluorescence from the total fluorescence. A script plotted background-subtracted fluorescence intensity over time for each well to allow visual inspection. If the dynamics of four positions looked similar, we randomly selected one to inspect. In rare occasions, all four positions were out-of-focus and were not used. In a small subset of experiments, a discontinuous jump in data appeared in all four positions for unknown reasons. We did not calculate rates across the jump. Occasionally, one or two positions deviated from the rest. This could be due to a number of reasons, including shift of focal plane, shift of field of view, black dust particles, or bright dust spots in the field of view. The outlier positions were excluded after inspecting the images for probable causes. If the dynamics of four positions differed due to cell growth heterogeneity at low concentrations of metabolites, all positions were retained.

We normalized total intensity against that at time zero, and averaged across positions. We calculated growth rate over three to four consecutive time points, and plotted the maximal net growth rate against metabolite concentration. If maximal growth rate occurred at the end of an experiment, then the experimental duration was too short and data were not used. For *L*^−^*H*^+^, the initial stage (3~4 hrs) residual growth was excluded from analysis due to residual growth supported by vacuolar lysine storage.

### Bioassay

75μl sample filtered through a 0.2μm filter was mixed with an equal volume of a master mix containing 2x SD (to provide fresh medium) as well as tester cells auxotrophic for the metabolite of interest (~1×10^4^ cells/ml, WY1340 over-night culture) in a flat-bottom 96-well plate. We then wrapped the plate with parafilm and allowed cells to grow to saturation at 30°C for 48 hrs. We re-suspended cells using a Thermo Scientific Teleshake (setting #5 for ~1 min) and read culture turbidity using a BioTek Synergy MX plate reader. Within each assay, SD supplemented with various known concentrations of metabolite were used to establish a standard curve that related metabolite concentration to final turbidity (e.g. Fig 3-Figure Supplement 4). From this standard curve, the metabolite concentration of an unknown sample could be inferred.

### Release assay

Detailed description of the release assay during lysine starvation can be found in (Hart et al., 2019a). Briefly, *L*^−^*H*^+^ strain was pre-grown in synthetic minimal media supplemented with high lysine (164 μM) to exponential phase. The cultures were washed in lysine-free media and allowed to starve for 2 hours at 30°C to deplete vacuolar lysine stores. Following starvation, the culture was periodically sampled (approximately every 6 hours for 24 hours) upon which live/dead cell densities were measured via flow cytometry (Methods “Flow Cytometry”), and culture samples were sterile filtered and supernatants were frozen. The supernatants were subjected to bioassay to measure hypoxanthine concentrations (Methods “Bioassay”). Hypoxanthine release rate can be inferred by the slope of the linear function relating integrated live cell density over time (cells/ml/hr) versus measured hypoxanthine concentration (μM). For an example, see Fig 3 – Figure Supplement 1.

To increase the throughput of this assay for screening chromosome truncation mutants (Fig 3B), we made the following modifications to this procedure. After measuring time zero cell densities by flow cytometry, we tracked fluorescence every 2 hours using automated 96-well plate fluorescence microscopy imaging (Hart et al., 2019b). We loaded 200 μL of OD~0.05 cells in SD per well while the rest of each culture was treated as in the normal assay to sterile filter at each sampling. Plate preparation, imaging, and images analysis were done as described in Methods: “Microscopy growth assay”. Fluorescence scales with live cell density, so we were able to estimate live cell densities at each time point *t* by taking (initial cell density)*(fluorescence intensity at time *t*)/(initial fluorescence intensity).

### Cell size measurements

Both Coulter counter and flow cytometry forward scattering can be used to compare the cell size distributions of yeast strains, with Coulter counter providing a direct measurement of cell size. We used the Z2 Coulter counter (Beckman), with the following settings: Gain=128; Current=0.5; Preamp Gain=224. We diluted cultures to OD_600_~0.01 to 0.3 (1 OD=7×10^7^ cells/ml) when necessary, sonicated cells (horn sonicator at low setting for 3 quick pulses or bath sonicator for 1 min), and placed 100 μl culture into Coulter cuvette. We then added 10ml sterilized isotone down the wall of the titled cuvette to avoid splashing, and analyzed the sample.

### Metabolite utilization in batch culture

We measured metabolite utilization after cells fully “saturated” the culture. We starved exponentially-growing cells (3-6 hours for *L*^−^*H*^+^, 24 hours for *H*^−^*L*^+^) to deplete initial intracellular stores and inoculated ~1×10^5^ cells/ml into various concentrations of the cognate metabolite up to 25 μM. We incubated for 48 hours and then measured cell densities by flow cytometry. We performed linear regression between input metabolite concentrations and final total cell densities within the linear range, forcing the regression line through origin. Utilization per birth in a saturated culture was quantified from 1/slope.

### Calculating uncertainty of ratio

Since release rate and metabolite utilization were measured in independent experiments, their errors were uncorrelated. For ratio *f* = *A/B*, suppose that *A* and *B* have standard deviations of *σ_A_* and *σ_B_*, respectively. Then *σ_f_* is calculated as 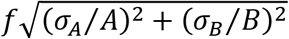.

### Chemostat

We have constructed an eight-vessel chemostat with a design modified from (Takahashi et al., 2015). For details of construction, modification, calibration, and operation, see (Skelding et al., 2017). A detailed discussion on using chemostats to quantify release and utilization phenotypes can be found in (Hart et al., 2019a). A summary is presented here.

For *L*^−^*H*^+^, due to rapid evolution, we tried to devise experiments so that live and dead populations quickly reached steady state. We first calculated the expected steady state cell density by dividing the concentration of lysine in the reservoir (20 μM) by fmole lysine utilized per new cell (Figure 4). We washed exponentially growing cells to remove extracellular lysine and inoculated 50%~75% of the volume at 1/3 of the expected steady state density. We filled the rest of the 19ml vessel with reservoir media (resulting in less than the full 20 μM of starting lysine, but more than enough for maximal initial growth rate, ~10-15 μM). We set the pump flow rate to achieve the desired doubling time *T* (19ml culture volume*ln(2)/T). We collected and weighed waste media for each individual culturing vessel to ensure that the flow rate was correct (i.e. total waste accumulated over time *t* was equal to the expected flow rate*t). We sampled cultures periodically to track population dynamics using flow cytometry (Methods, “Flow cytometry”), and filtered supernatant through a 0.45 μm nitrocellulose filter and froze the supernatant for metabolite quantification at the conclusion of an experiment (Methods, “Bioassay”). At the conclusion of an experiment, we also tested input media for each individual culturing vessel to ensure sterility by plating a 300 μl aliquot on an YPD plate and checking for growth after two days of growth at 30°C. If a substantial number of colonies grew (>5 colonies), the input line was considered contaminated and data from that vessel was not used. For most experiments, we isolated colonies from end time point and checked percent evolved (Methods, “Detecting evolved clones”). For *L*^−^*H*^+^, we only analyzed time courses where >90% of population remained ancestral.

In a lysine-limited chemostat, live cell density [*L*^−^*H*^+^]_*live*_ is increased by growth (at a rate *g*), and decreased by dilution (at a rate *dil*):

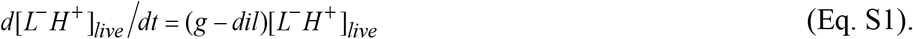

*L*, lysine concentration in the culturing vessel, is increased by the supply of fresh medium (at concentration *L_0_*), and decreased by dilution and utilization (with birth of each new cell utilizing *u_L_* amount of lysine).

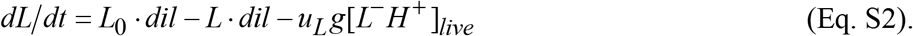

Finally, hypoxanthine concentration *H* is increased by release (from live cells at *r_H_* per live cell per hr, (Hart et al., 2019a)), and decreased by dilution.

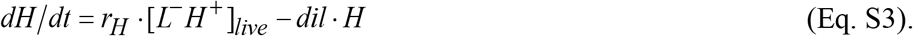

Note that at the steady state (denoted by subscript “ss”), growth rate is equal to dilution rate (setting Eq. 4 to zero):

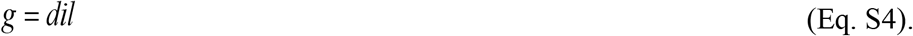

To measure metabolite utilized per birth at steady state, we set Eq. S2 to zero

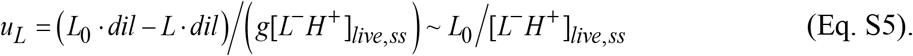

Here, the approximation holds because the concentration of lysine in chemostat (*L*) is much smaller than that in reservoir (*L_0_*).

To measure release rate at steady state, we can set Eq. S3 to zero and obtain:

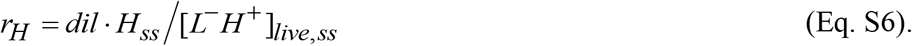

Thus, the exchange ratio can be quantified from

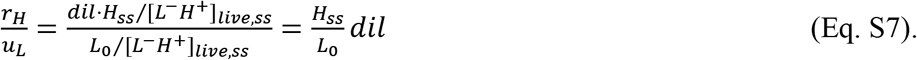

### Community growth rate

This derivation is adapted from (Hart et al., 2019a). If we culture *L*^−^*H*^+^ with *H*^−^*L*^+^, as the case in a spatially–structured environment, we have

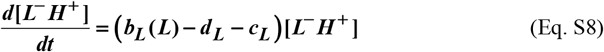

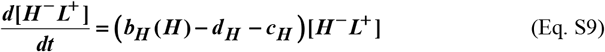

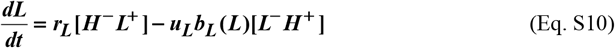

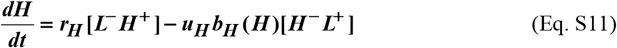

Equation S8 states that the clonal population density *L*^−^*H*^+^ increases at birth rate *b_L_* which in turn depends on the concentration of lysine *L*, and decreases at death rate *d_L_* and cost of metabolite overproduction *c_L_*. Equation S9 describes how clonal population density *H*^−^*L*^+^ changes over time. Equation S10 states that the concentration of lysine *L* increases due to releaser *H*^−^*L*^+^ releasing at a rate *r_L_* and decreases as *u_L_* amount is utilized per birth of consumer *L*^−^ *H*^+^. Equation S11 describes how the concentration of *H* changes over time. All parameters are non-negative. Note that only a single genotype per species is considered.

We can calculate the steady state growth rate *g_comm_*. Since strain ratio becomes fixed, both strains must grow at the same rate as the community. This also means that *L* and *H* concentrations do not change.

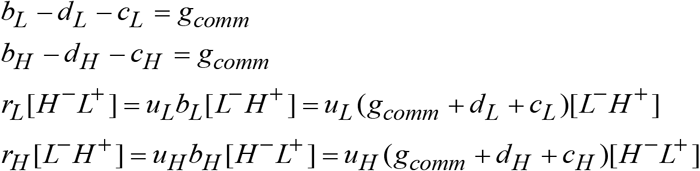

Multiply the last two equations, we get

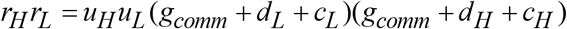

Solving this, we get

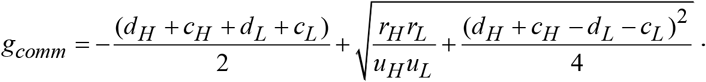

When *d_H_* and *d_L_* and *c_H_* and *c_L_* are small compared to 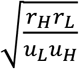, which is the case for CoSMO, we have

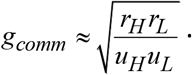

### Quantifying spatial growth dynamics

Briefly, exponentially growing *L*^−^*H*^+^ and *H*^−^*L*^+^ were washed free of lysine and hypoxanthine supplements, respectively. *H*^−^*L*^+^ cells were further starved for 24 hrs to reduce CoSMO growth lag phase (Hart et al., 2019a). The two strains were then mixed at approximately 1:1 ratio, and 15 μl of 4×10^4^ total cells were spotted on the center of agarose pads (1/6 of a petri dish pie), forming an inoculum spot of radius ~4 mm. Periodically, cells from pads were washed off into water and subjected to flow cytometry. The agarose pad generally contained 0.7 μM lysine, although including or not including this low concentration of lysine did not make a difference in the steady-state community growth rate.

---

**Fig 1 - Figure Supplement 1.**
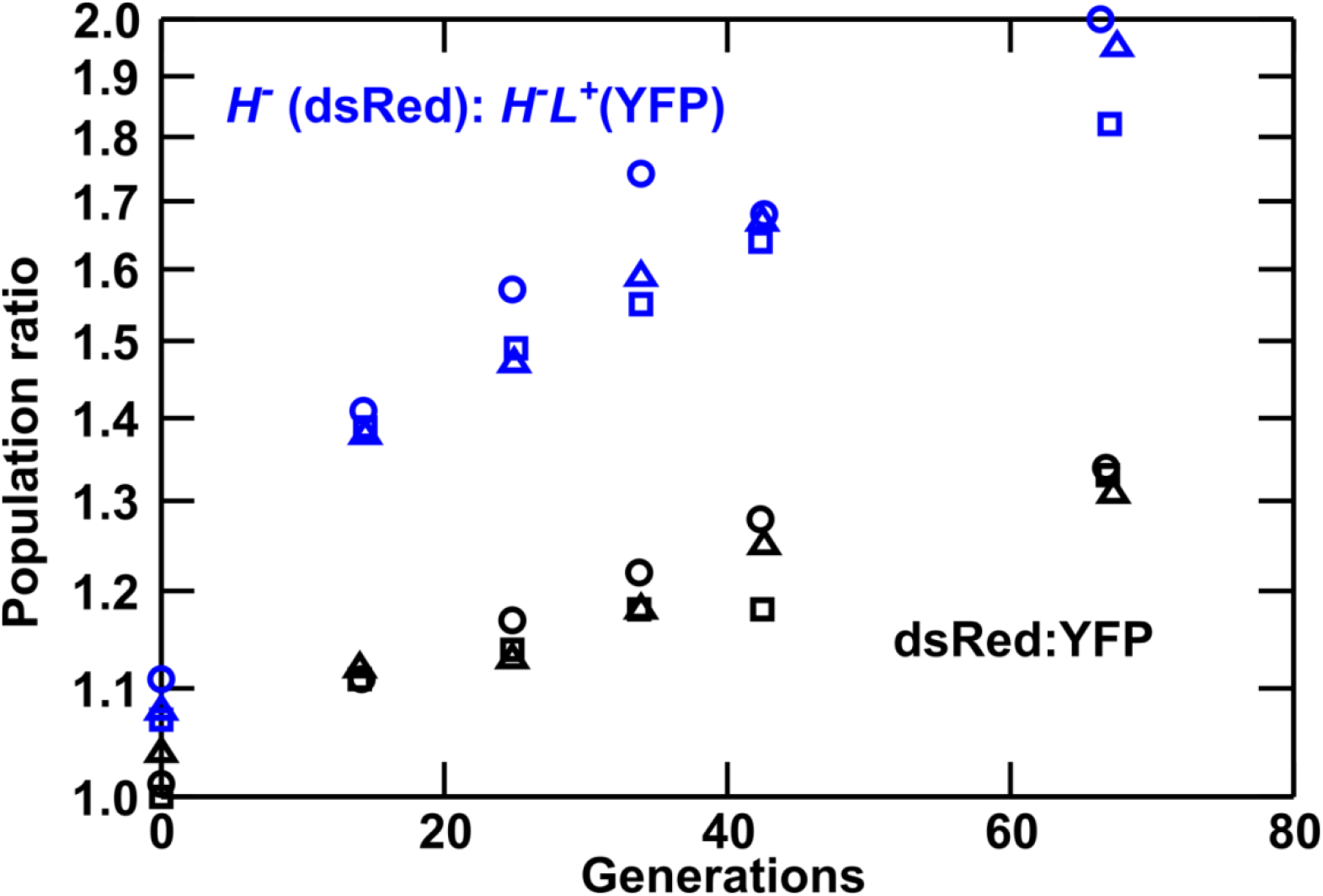
Lysine overproduction incurs a fitness cost. Exponentially-growing isogenic strain pairs were competed in triplicate cocultures (circle, square, and triangle) at 30°C in minimal SD medium with adenine supplemented in excess (108.6 μM) when necessary. Cocultures were diluted twice daily to avoid depletion of nutrients. Population ratios were monitored using flow cytometry. In blue curves, dsRed-marked *H*^−^ not overproducing lysine (WY944) has a fitness advantage of 1.2+/-0.2% over YFP-marked *H*^−^*L*^+^ overproducing lysine (WY954). In black curves, dsRed-marked WY926 has a fitness advantage of 0.6+/-0.1% over YFP-marked WY923. Thus, lysine overproduction imposes a fitness cost of ~0.6%. Data can be found in Fig 1-Figure Supplement 1 Source Data.

**Fig 2 - Figure Supplement 1.**
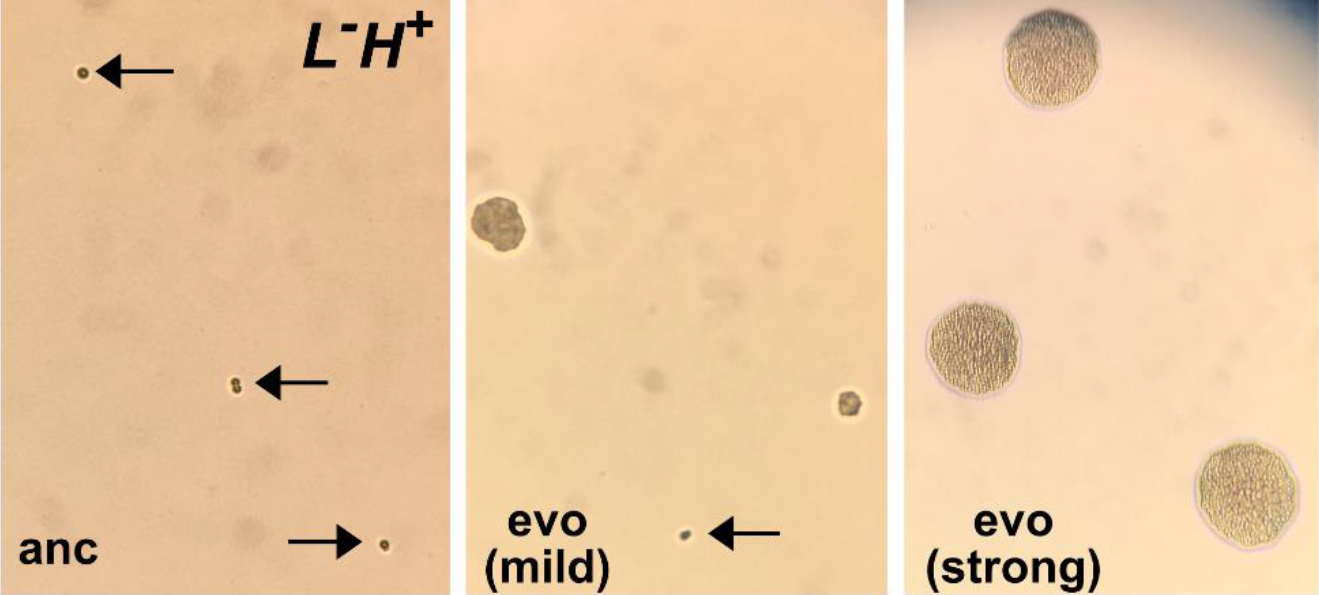
Micro-colony assay reveals improved growth in low lysine. A visual assay that distinguishes ancestral from evolved *L*^−^*H*^+^ clones. An ancestral clone (left) and two evolved *L*^−^*H*^+^ clones (center and right) were plated on SD plates supplemented with 1.5 μM lysine. Ancestral cells (WY1335, left) failed to divide (arrows). Cells from a mildly-adapted evolved clone (harboring duplication of Chromosome 14, center) showed heterogeneous phenotypes: some cells remained undivided (arrow), while other cells formed microcolonies of various sizes. Cells from a strongly-adapted evolved clone (harboring an *ecm21* mutation, right) formed microcolonies of a uniform and large size. These images were taken using a cell phone camera and thus do not have a scale bar. For reference, an average yeast cell has a diameter of ~5 μm. This figure and figure legend are adapted from (Hart et al., 2019a).

**Fig 2 - Figure Supplement 2.**
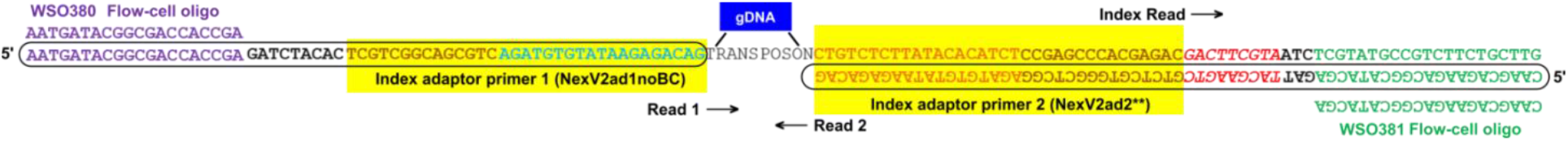
Whole genome sequencing Nextera V2. Schematic for whole genome sequencing using Nextera V2. Sequences for primers can be found in Table S3. The cyan and orange regions right next to gDNA are added by the transposase during DNA fragmentation (tagmentation).

**Fig 3 - Figure Supplement 1.**
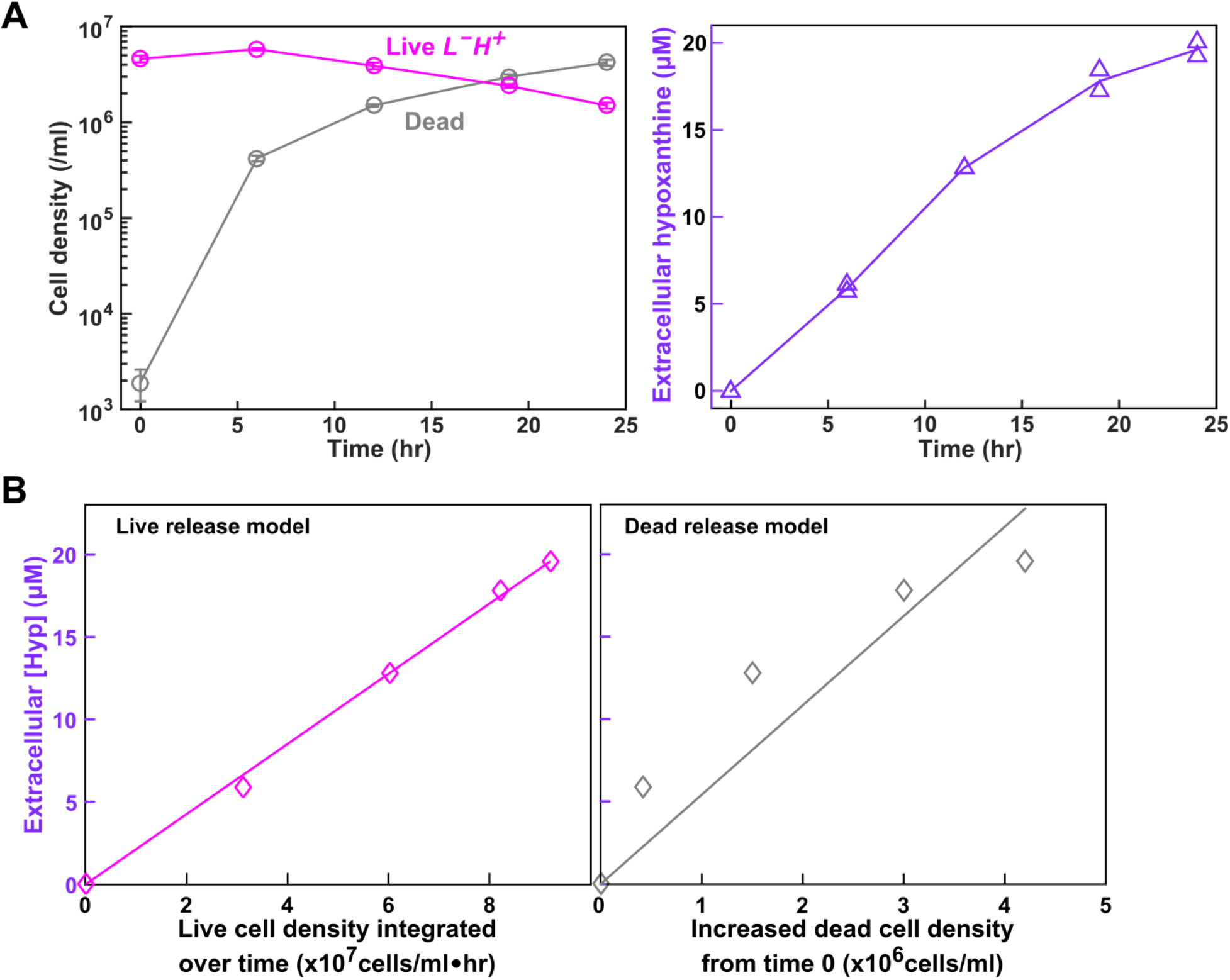
Starvation batch culture release assay. This figure is adapted from Figure 5 of (Hart et al., 2019a). (**A**) Hypoxanthine release during lysine starvation. Exponential *L*^−^*H*^+^ cells were washed free of lysine, and diluted into SD without lysine. Live and dead population densities and supernatant hypoxanthine concentrations were measured over time (Methods, “Flow cytometry”; “Bioassay”). (**B**) (Left) If hypoxanthine was released by live cells at a constant rate, then hypoxanthine concentration should scale linearly against live cell density integrated over time. (Right) If hypoxanthine was released upon cell death, then hypoxanthine concentration should scale linearly against dead cell density. Live release model has better linearity than dead release model, and therefore hypoxanthine is likely released by live cells. The conclusion of live release was corroborated by metabolite extraction experiments (Hart et al., 2019a). Data used to generate these plots can be found in (Hart et al., 2019a).

**Fig 3 - Figure Supplement 2.**
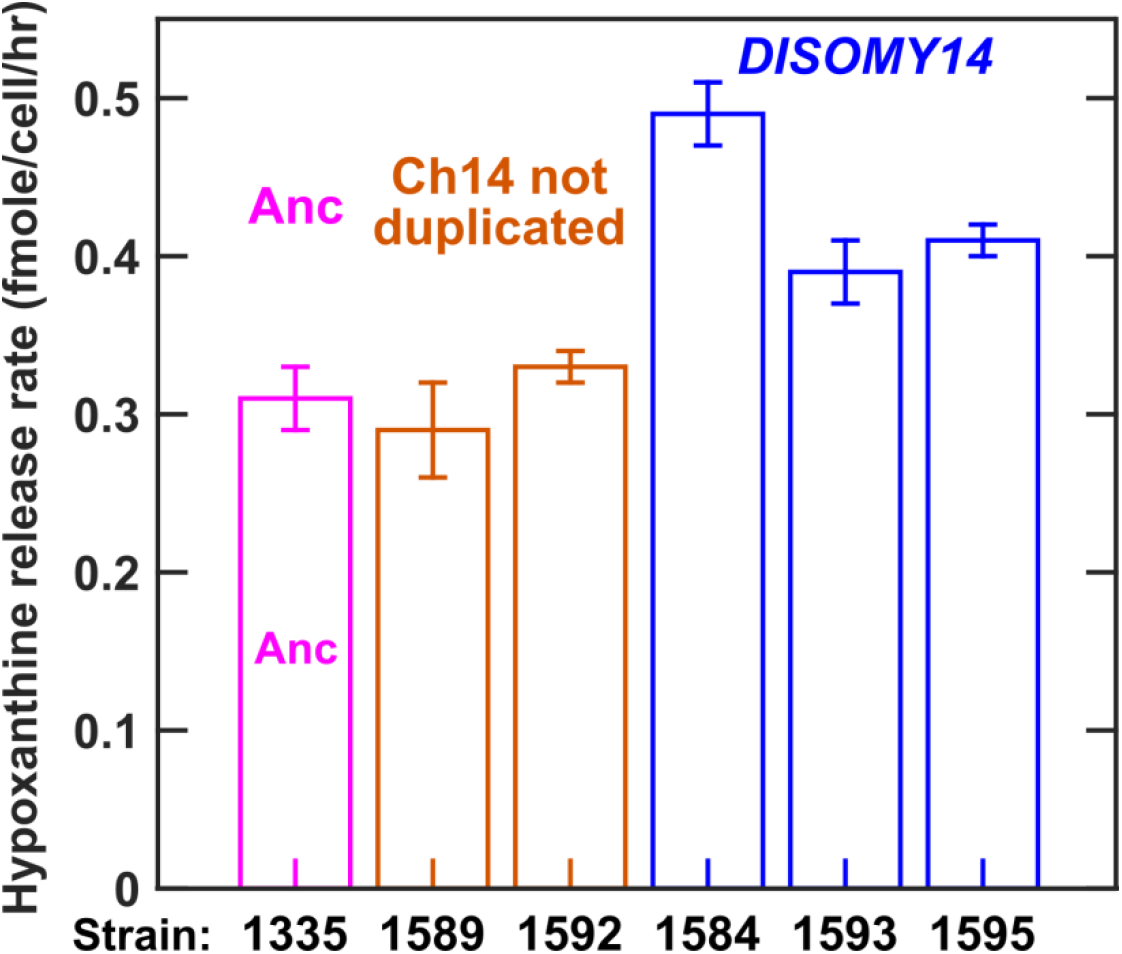
Evolved L^−^H^+^ clones with improved hypoxanthine release rates harbored Chromosome 14 duplication. Our previous work has established that hypoxanthine is released by live *L*^−^*H*^+^ cells (Hart et al., 2019a)), and that hypoxanthine release rate is relatively constant regardless of whether cells are starved of lysine or grown in lysine-limited chemostats mimicking the community environment (Hart et al., 2019a). Since starvation release is easier to measure, we quantified hypoxanthine release rates of *L*^−^*H*^+^ clones in starvation batch cultures (Methods, “Release assay”). As shown later, we obtained similar results in lysine-limited chemostats mimicking the CoSMO environment. Specifically in this experiment, we grew ancestral and evolved *L*^−^*H*^+^ clones to exponential phase in SD supplemented with non-limiting lysine, and washed the cultures free of lysine with SD. We then starved each culture for 24 hours at 30°C during which we sampled approximately every 6 hours to measure live cell density (Methods, “Flow cytometry”) and supernatant hypoxanthine concentration (Methods, “Bioassay”). From these dynamics, we calculated hypoxanthine release rate (Methods, “Starvation release assay”; Fig 3-Figure Supplement 1). Release rates are plotted here with error bars indicating 2 standard errors of mean (SEM) of at least four measurements. Evolved clones harboring chromosome 14 duplication (blue) exhibited increased hypoxanthine release rate per cell in comparison to the ancestor (magenta), while evolved clones without chromosome 14 duplication (orange) exhibited the ancestral release rate. Data can be found in Fig 3-Figure Supplement 2 Source Data.

**Fig 3 - Figure Supplement 3.**
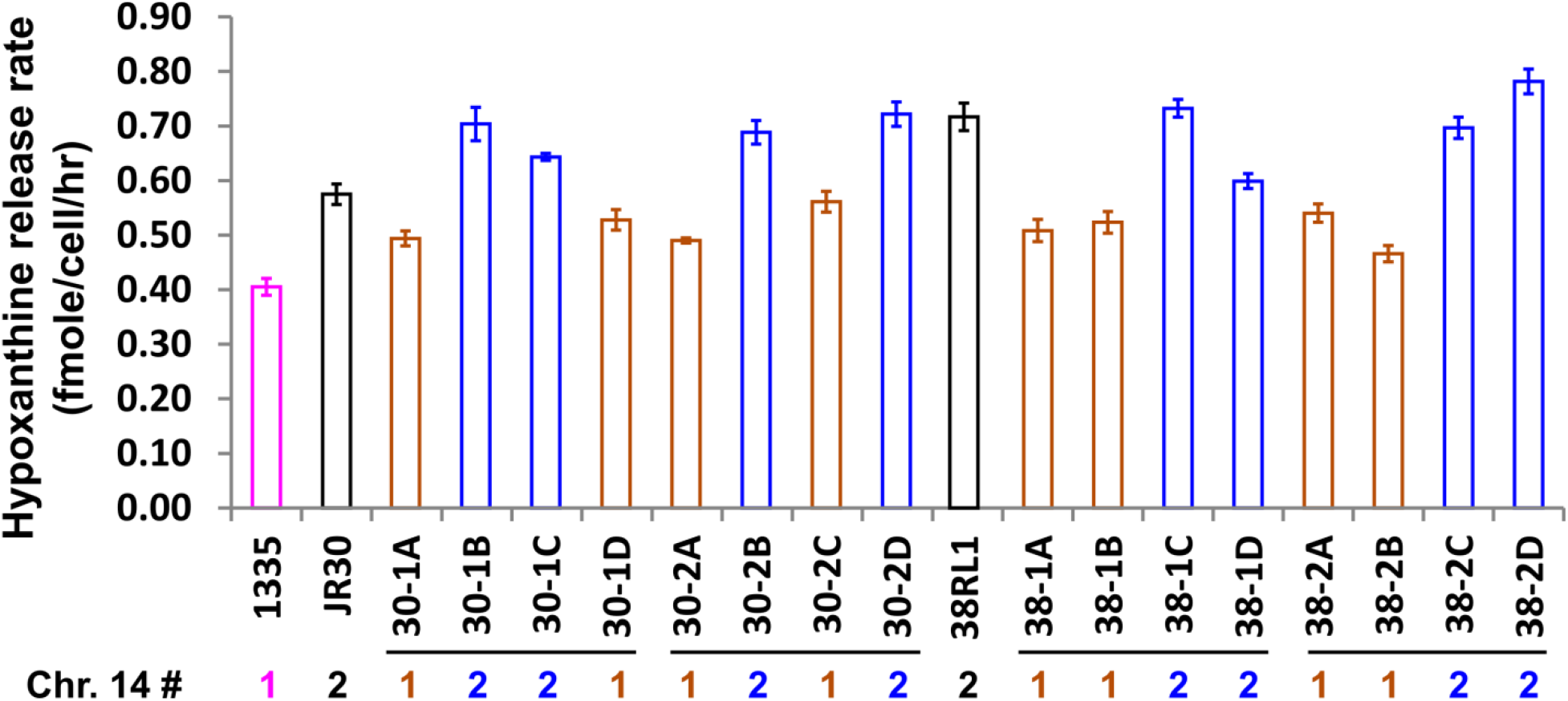
Increased release rate segregates with DISOMY14. We crossed evolved clones JR30 (WY1593; black) and 38RL1 (WY1584; black) containing *DISOMY14* with an ancestral strain containing one copy of chromosome 14 (WY1335; magenta). When we dissected tetrads (4 spores/tetrad as marked by horizontal lines), the hypoxanthine over-release phenotype (blue, measured in starvation batch cultures) segregated with *DISOMY14* (Methods, “RADseq”). Data can be found in Fig 3-Figure Supplement 3 Source Data, Sheet “Plots”.

**Fig 3–Figure Supplement 4.**
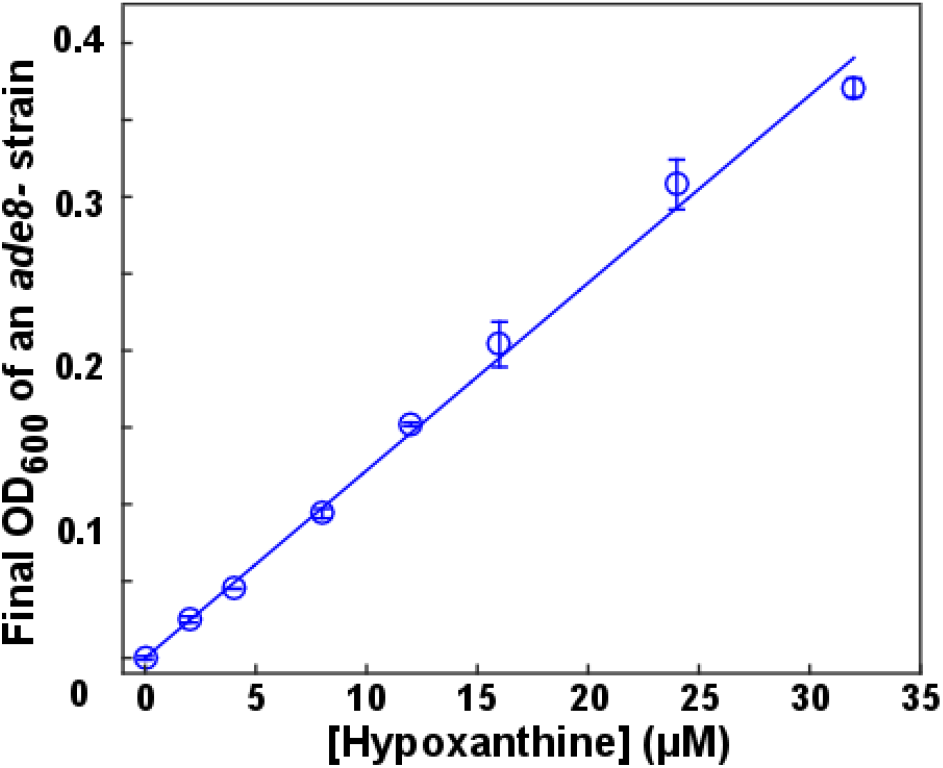
Bioassay. This figure is adapted from S3 Fig of (Hart et al., 2019a). The final turbidity of an *ade8*-(WY1340) tester strain increases with increasing concentrations of hypoxanthine (blue) in a linear fashion (within the tested concentration range). Data can be found in (Hart et al., 2019a).

**Fig 3 - Figure Supplement 5.**
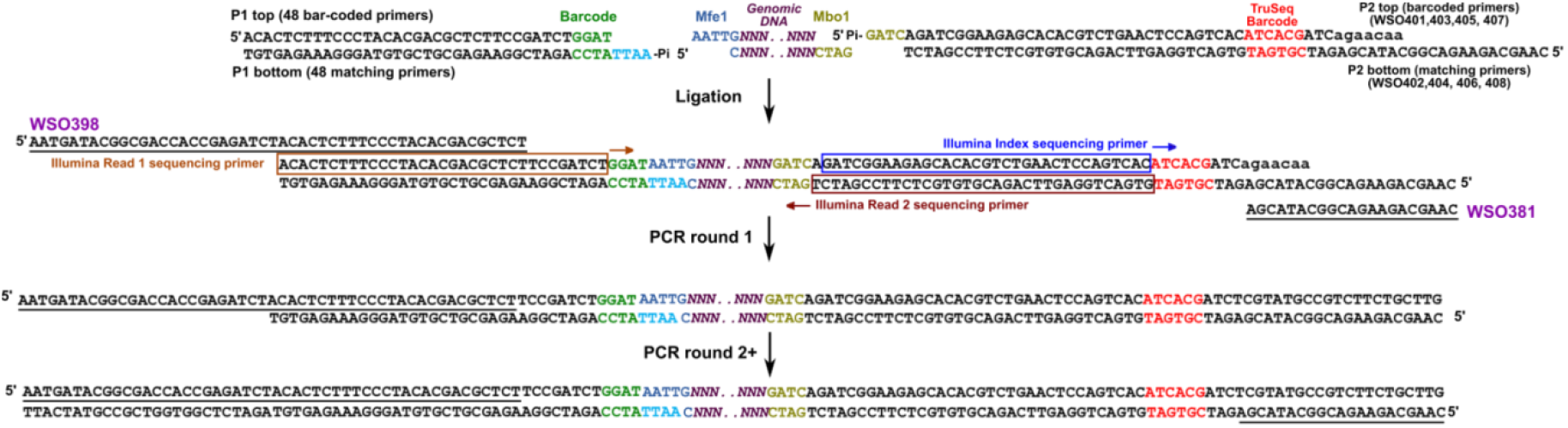
RADseq. Schematic for RADseq using TruSeq. For detailed information, see Methods “RADseq”. Sequences for primers can be found in Table S4.

**Fig 4–Figure Supplement 1.**
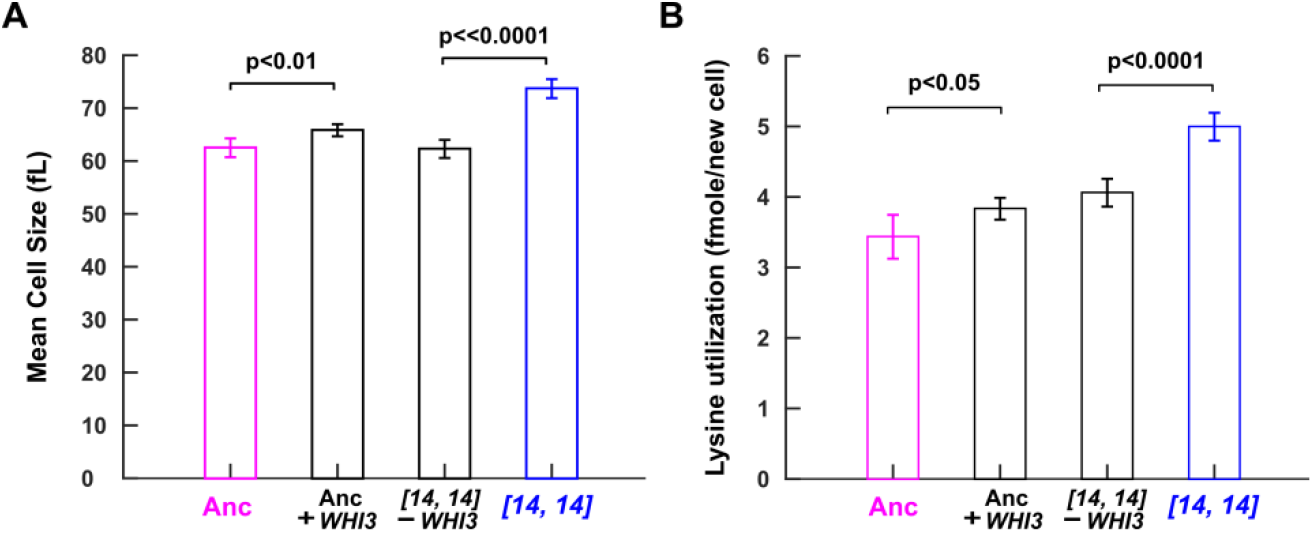
WHI3 duplication causes increased cell size and increased lysine utilization per birth. Introducing an extra copy of *WHI3* into the ancestral background (WY2357~2359) increased cell size and lysine utilization per birth. Deleting the duplicated *WHI3* from *DISOMY14* (WY2350~2352) decreased cell size and lysine utilization per birth. Duplicating or deleting *WHI3* did not always result in a full phenotype switch to or from *DISOMY14* phenotype, but always resulted in a significant change in phenotype from the parent strain. For example, duplicating *WHI3* in the ancestor resulted in a significant increase in lysine required per new cell, but not to the level of *DISOMY14*. This could be due to the integrated *WHI3* copy not being expressed to the same extent as the duplicated *WHI3* copy in *DISOMY14*, or due to duplication of other genes in *DISOMY14*, or due to mutations in other chromosomes of the *DISOMY14* strain. Mean and two SEM are plotted. P-values are from two-tailed t-test assuming equal variance (equal variance being tested via the F-test). A and B were measured in batch cultures. Data for A and B can be found in Fig 4 Source Data, “CellSize” sheet and “Consumption” sheet, respectively.

**Fig 4 - Figure Supplement 2.**
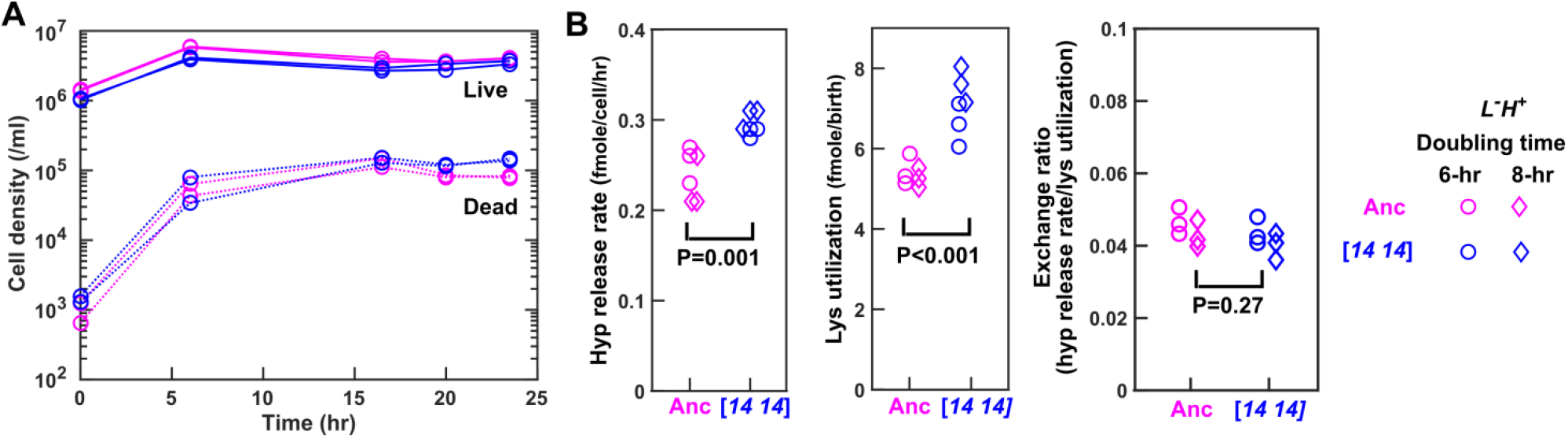
In chemostats, DISOMY14 and ancestor share identical exchange ratio. (**A**) Chemostat population dynamics. Ancestor (WY1335, magenta) or *DISOMY14* (WY2349, blue) *L*^−^*H*^+^ cells were grown in separate, lysine-limited chemostats (Methods “chemostats”). Live and dead population densities were tracked using flow cytometry (Methods, “Flow cytometry”). Supernatant hypoxanthine concentrations are plotted in Fig 4D. (**B**) *DISOMY14* is not partner-serving. We quantified hypoxanthine release rate (Methods “chemostats” Eq. S6), lysine utilization per new cell (Methods “chemostats” Eq. S5), and exchange ratio (Methods “chemostats” Eq. S7) for ancestor and *DISOMY14*, using the steady state data from 15~24 hrs. Compared to the ancestor, *DISOMY14* displayed higher hypoxanthine release rate per cell and higher lysine utilization per birth, regardless of whether we treated 6-hr doubling time chemostat data (circles) and 8-hr doubling time chemostat data (diamonds) as separate groups or as a single group. P values were calculated from two-tailed *t*-test assuming equal variance (equal variance being tested by the F-test). Ancestor and *DISOMY14* have significantly different hypoxanthine release rate per cell and lysine utilization per birth, but their exchange ratios are similar. Data can be found in Fig 4 Source Data.

**Fig 4 – Figure Supplement 3.**
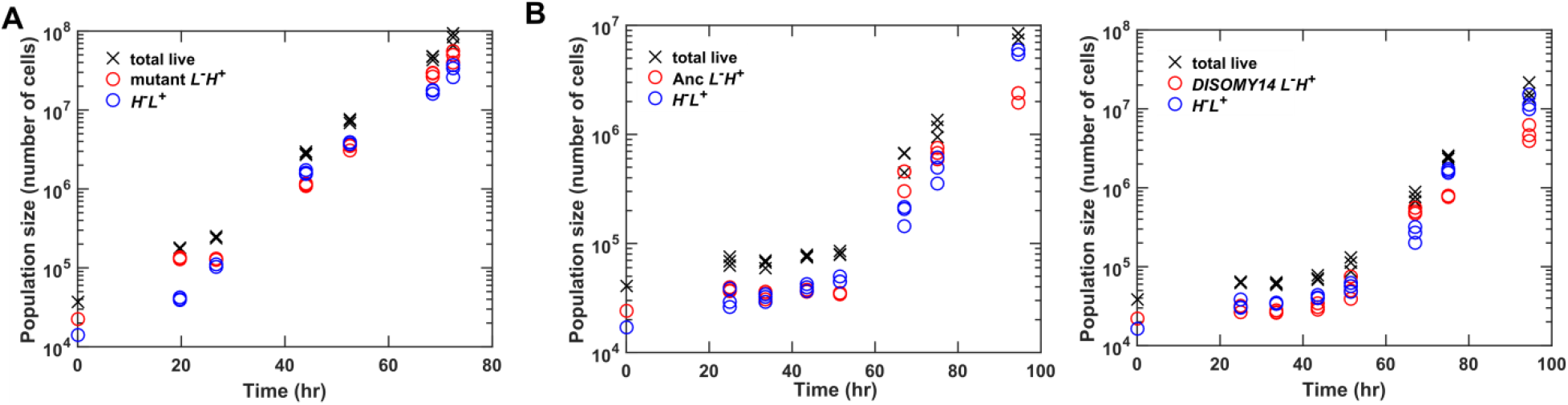
CoSMO growth dynamics. (**A**) Exponential growth lasting to 10^8^ total cells. (**B**) Examples of CoSMO dynamics consisting of ancestral or *DISOMY14 L*^−^ *H*^+^. The duration of the lag phase can vary with the size of the inoculation spot (and the initial surface cell density). Data can be found in Fig 4 Source Data, “Dynamics”.

***Table S1. Strain table***

Table S1.

***Table S2. Mutations in evolved L^−^H^+^***

Table S2

***Table S3. Whole genome sequencing primer sequences***

Table S3

***Table S4. RADseq primer sequences***

Table S4

## Acknowledgement

We thank Gareth Cromie and Eric Jeffery in Aimee Dudley lab for advice on RADSeq and for providing reagents, Dan Gottschling for WSB174 (pADH4UCA-IV) plasmid, Sergey Kryazhimskiy for whole genome sequencing library preparation protocol, Klavins lab for Gibson assembly protocol, and Jacob Kitzman from Jay Shendure lab for bar-code index sequences. We thank Li Xie, Alex Yuan, and David Skelding for discussions, Maxine Linial and Delia Pinto-Santini for critical reading of the initial manuscript, and Kevin Foster (University of Oxford) and Ronald Noë for critical reading of the revised manuscript.

